# Enhancing structural heterogeneity in managed forest landscapes promotes gamma but not beta diversity in understorey plant communities

**DOI:** 10.1101/2025.08.29.673108

**Authors:** Pia M. Bradler, Benjamin M. Delory, Sebastian Dittrich, Christian Ammer, Claus Bässler, Marc W. Cadotte, Anne Chao, Po-Yen Chuang, Werner Härdtle, Oliver Mitesser, Akira Mori, Jörg Müller, Keita Nishizawa, Fons van der Plas, Goddert von Oheimb, Andreas Fichtner

**Affiliations:** Institute of Ecology, Leuphana University of Lüneburg, Lüneburg, Germany; Institute of General Ecology and Environmental Protection, TUD Dresden University of Technology, Tharandt, Germany; Copernicus institute of sustainable development, Utrecht University, The Netherlands; Department Silviculture and Forest Ecology of the Temperate Zones, University of Goettingen, Germany; Fungal Ecology, University of Bayreuth, Universitätsstraße 30, Bayreuth, Germany; Bavarian Forest Nationalpark; Department of Biological Sciences, University of Toronto Scarborough, Toronto, ON, Canada; Institute of Statistics, National Tsing Hua University, Hsinchu, Taiwan; Chair of Conservation Biology and Forest Ecology, Biocenter of the Julius-Maximilians University of Würzburg; Research Center for Advanced Science and Technology, The University of Tokyo, Tokyo, Japan; Plant Ecology and Nature Conservation Group, Wageningen, The Netherlands

**Keywords:** Biodiversity, diversity facets, forest structure, heterogeneity–diversity relationships, Hill-Chao numbers, sample coverage standardisation, spatial scales, structural heterogeneity, understorey vegetation

## Abstract

1. Although habitat heterogeneity is known to enhance local species diversity, the effects of management-driven structural heterogeneity on understorey plant communities across spatial scales remain poorly understood, despite their crucial role for forest biodiversity and ecosystem functioning.
2. To analyse how forest understorey plant communities respond to an enhancement of structural heterogeneity in managed forests, we established 11 experimental sites consisting of two paired forest landscapes, an untreated homogenous control and a treatment district (ESBC). In treatment districts, structural heterogeneity was enhanced through different combinations of local patch-scale manipulations of light and deadwood features, leading to greater between-patch heterogeneity at the landscape scale. We performed a meta-analysis across these 11 sites using a Hill-Chao number and sample coverage standardisation framework.
3. Gamma diversity increased across taxonomic, functional and phylogenetic facets in structurally heterogeneous forests (ESBC districts) via higher alpha diversity. This effect was positively associated with heterogeneity in light availability between forest patches, but not with their mean light availability. In contrast, we found no support that species turnover among patches (i.e., beta diversity) significantly contributes on average to the observed increase in gamma diversity. However, both the direction and magnitude of beta diversity responses varied substantially among landscapes.
4. On average, structurally heterogeneous forests supported higher species richness for both open and closed forest habitat species.
5. *Synthesis and applications:* Our findings highlight the benefits of enhancing structural heterogeneity for understorey plant diversity in managed forest landscapes. Specifically, management strategies that create a spatial mosaic of interventions, such as combining single-tree removal with group felling, can increase the variety of light niches among forest patches, thereby supporting the conservation of a wide range of understorey plant species, including forest specialists.

## 1. Introduction

Biodiversity is increasingly threatened by intense human pressures, particularly through habitat changes, fragmentation and losses (Keck et al. 2025; Brooks et al. 2002). These pressures are driving widespread biodiversity loss and population declines that can decrease the dissimilarity in species composition (beta diversity) between local communities, i.e. biotic homogenisation (Dornelas et al. 2014; Gossner et al. 2016). For Central European forests, a shift towards a dominance of shady and nutrient-rich habitat conditions has been observed (Vild et al. 2024), resulting in changes in the diversity and composition of understorey plant communities. For example, it has been shown that understorey vegetation is undergoing biotic homogenisation, which is characterised by the replacement of small-ranged species by nutrient-demanding and shade-tolerant species often associated with closed-canopy conditions (Dirnböck et al. 2014; Sanczuk et al. 2024; Staude et al. 2020). Environmental heterogeneity, characterised by a diversity of habitats, can sustain distinct species assemblages at larger spatial scales, thereby maintaining higher beta and gamma diversity and counteracting biotic homogenisation. While enhancing heterogeneity could be a goal of habitat management (Müller et al. 2023), species losses and gains occur differently across spatial scales and how patterns observed at the local (alpha) diversity scale translate into changes at the regional (gamma) diversity scale is not fully understood (McGill et al. 2015).

Following the habitat heterogeneity hypothesis (MacArthur and MacArthur 1961), an increase in the spatial heterogeneity in habitat conditions is directly linked to an increase in species diversity, as more available niche space allows more species to coexist. In a meta-analysis, Stein et al. (2014) found strong quantitative evidence for a general positive relationship between environmental heterogeneity and species richness across many taxa and in different regions worldwide. However, a recent study in temperate forests showed that the shape of the heterogeneity–diversity relationship strongly depends on species groups (e.g. plants, fungi, lichens, birds, arthropods; Heidrich et al. 2020). Additionally, heterogeneity–diversity relationships may also be shaped by different diversity facets, such as taxonomic, functional and phylogenetic diversity (Cadotte et al. 2013). For example, de Pauw et al. (2021) showed that taxonomic diversity of herbaceous understorey plant species increased with increasing light availability, while functional diversity increased with stronger microclimate buffering (i.e., in dense stands). Conversely, phylogenetic diversity was not affected by changes in light availability in this study, but increased with increasing soil acidity. Consequently, focusing solely on species richness could mask changes in species abundance, and therefore the effects of increased heterogeneity on community structure (Boch et al. 2013; Cramer & Willig, 2005; Dornelas et al. 2009; Hedwall and Brunet 2016). Moreover, rare species with small population sizes and high degrees of specialisation are likely to be disproportionately affected by changes in habitat conditions (Dornelas et al. 2009; Staude et al. 2020). Analysing how rare species, species with different environmental preferences (e.g. forest specialists vs. generalists), and functionally or phylogenetically distant species respond to habitat changes is therefore crucial to better understand community dynamics and biodiversity responses to environmental changes.

Forest understorey plant communities represent a major component of temperate forest biodiversity, harbouring on average more than 80% of vascular plant species (Gilliam 2007). Moreover, they have a strong impact on tree recruitment, provide habitats and floral resources for higher trophic levels, and play an important role for nutrient cycling (Landuyt et al. 2019). Forest understorey plants are also known to quickly respond to changes in habitat conditions and resource availability, respectively, with many species being specialised to specific light conditions, tree species and microclimates associated with closed or open canopies (von Oheimb and Härdtle, 2009; Heidrich et al. 2020; de Frenne et al. 2019). Changes in canopy structure are therefore expected to alter the diversity and composition of understorey plant communities (De Frenne 2024). Several studies have shown that local modifications of the canopy density can increase understorey plant diversity due to higher light availability (Bernhardt-Römermann et al. 2015; De Frenne et al. 2013; Dormann et al. 2020). Other studies, however, suggest that heterogeneous light conditions are more important for regulating understorey plant diversity than the amount of light *per se* (Helbach et al. 2022; Tinya & Ódor, 2016). In addition, canopy openings not only affect light conditions, but also wind speed, air humidity, soil temperature and moisture, litter input and thus nutrient availability at the forest floor (Ritter et al. 2005; Abd Latif & Blackburn, 2010).

Managed forests in their optimal timber production phase are often characterised by relatively homogenous low-light conditions and low understorey plant diversity. This is largely attributable to the absence of early and late successional stages, which are typically associated with more open canopy structures and a high diversity of various taxa (Hilmers et al. 2018). At the landscape scale, the availability of habitat patches of different successional stages is assumed to be a key driver for promoting understorey plant diversity (Schall et al. 2018; Hylander et al. 2022; Duflot et al. 2022). In managed forests, environmental heterogeneity can be enhanced through management interventions, such as manipulating light and deadwood availability (Heidrich et al. 2020). Evidence for the effects of structural heterogeneity on biodiversity, however, is limited to observational or modelling studies (see e.g. van der Plas et al. 2016; Schall et al. 2020).

Here, we analyse how understorey plant communities respond to experimentally enhanced structural heterogeneity across multiple spatial scales, using data from 234 patches nested within 11 replicated experimental sites in temperate managed forests across Germany (BETA-FOR experiment). Each experimental site consisted of two paired forest districts, a control and a treatment district (Müller et al. 2023). In each treatment district, structural heterogeneity was enhanced through different combinations of local-scale manipulations of light conditions and deadwood characteristics (Enhancement of Structural Complexity, ESC *sensu* Keeton 2006), resulting in increased structural heterogeneity at the landscape scale (Enhancement of Structural Beta-Complexity, ESBC). In contrast, control districts were not subjected to any intervention aimed at enhancing stand structural complexity. This experimental design allowed us to quantify changes in various diversity facets (taxonomic, functional, and phylogenetic) of understorey plant communities in response to enhanced structural heterogeneity across spatial scales (alpha, beta and gamma). Specifically, we tested the following hypotheses:

H1: Following the habitat heterogeneity hypothesis, we expect landscapes with greater structural heterogeneity to support higher gamma diversity of understorey plants.
H2: Gamma diversity is primarily driven by species turnover (beta diversity), rather than by increased local understorey plant diversity (alpha diversity).
H3: Increased structural heterogeneity at the landscape scale enhances habitat suitability and niche availability by creating spatial variation in light conditions, thereby promoting higher species richness among both forest specialists and generalists.

## Material and methods

### 2.1 Study sites and experimental design

We established 11 experimental sites in temperate managed forests across Germany covering a wide range of different biotic and abiotic conditions. Each site consisted of two paired forest districts, a control and a treatment district (Müller et al. 2023). In the treatment district (ESBC), we enhanced structural heterogeneity by manipulating light conditions and deadwood features at the local patch scale (ESC, Enhancement of Structural Complexity), creating a spatial mosaic of differently treated neighbouring patches. No tree removal or deadwood manipulation was applied in the control districts. Interventions to increase structural heterogeneity were implemented in treatment districts between the winters of 2015/2016 and 2018/2019 in the different sites. The districts were established in predominantly beech-dominated (*Fagus sylvatica* L.) forest stands that span an elevational gradient from 35 m to 1055 m above sea level.

In ESBC districts, greater spatial variation in light conditions was achieved through either spatially aggregated (gap felling) or distributed (selective thinning) tree removal of ∼30% of trees within a forest patch. Manipulations to increase deadwood features included leaving snags, logs, stumps or a combination of snags and logs after tree removal. These interventions jointly altered the availability of two key ecological resources in forests: light and deadwood (Müller et al. 2023). Each patch was subjected to a specific factorial combination of canopy structure and deadwood manipulation, resulting in eight structurally distinct treatment patches per district. To further enhance structural heterogeneity among patches, one additional untreated control patch was included in each district, resulting in a total of nine patches per district. In three experimental landscapes (U01, U02, U03), six additional treatments were implemented in ESBC districts, resulting in 15 patches per district. This experimental design enhanced structural complexity among patches—i.e., structural beta-complexity—thereby allowing analysis of diversity pattern at multiple spatial scales: alpha (within-patch, n=234 across all sites), beta (among-patch) and gamma scale (landscape-scale), across 11 experimental landscapes.

### 2.2 Vegetation data

All 234 patches were surveyed in summer 2023. Plant species occurrence and cover was recorded in five circular sub-patches per patch, each with a radius of 4 m (50 m^2^) (Figure 1). Within each patch, the sub-patches were arranged with one in the centre and the remaining four positioned at the corners. These corner sub-patches were located at a distance of 12 m (six study sites) or 21 m (5 study sites; owing to the design of a previous study) from the centre of the patch. We recorded all vascular plants including woody species (tree or shrub seedlings and juveniles) up to a height of 1 m in vegetation surveys (c.f. Gilliam 2007). In total, we recorded 226 plant species. Species names were standardised using the Taxonomic Name Resolution Service 0.3.4 (TNRS R package, Maitner and Boyle 2023). Two patches had to be removed from the dataset due to requirements of our statistical approach as no species were recorded in them, resulting in only eight patches in B06 (control district) and B07 (control district).

**Figure 1.**
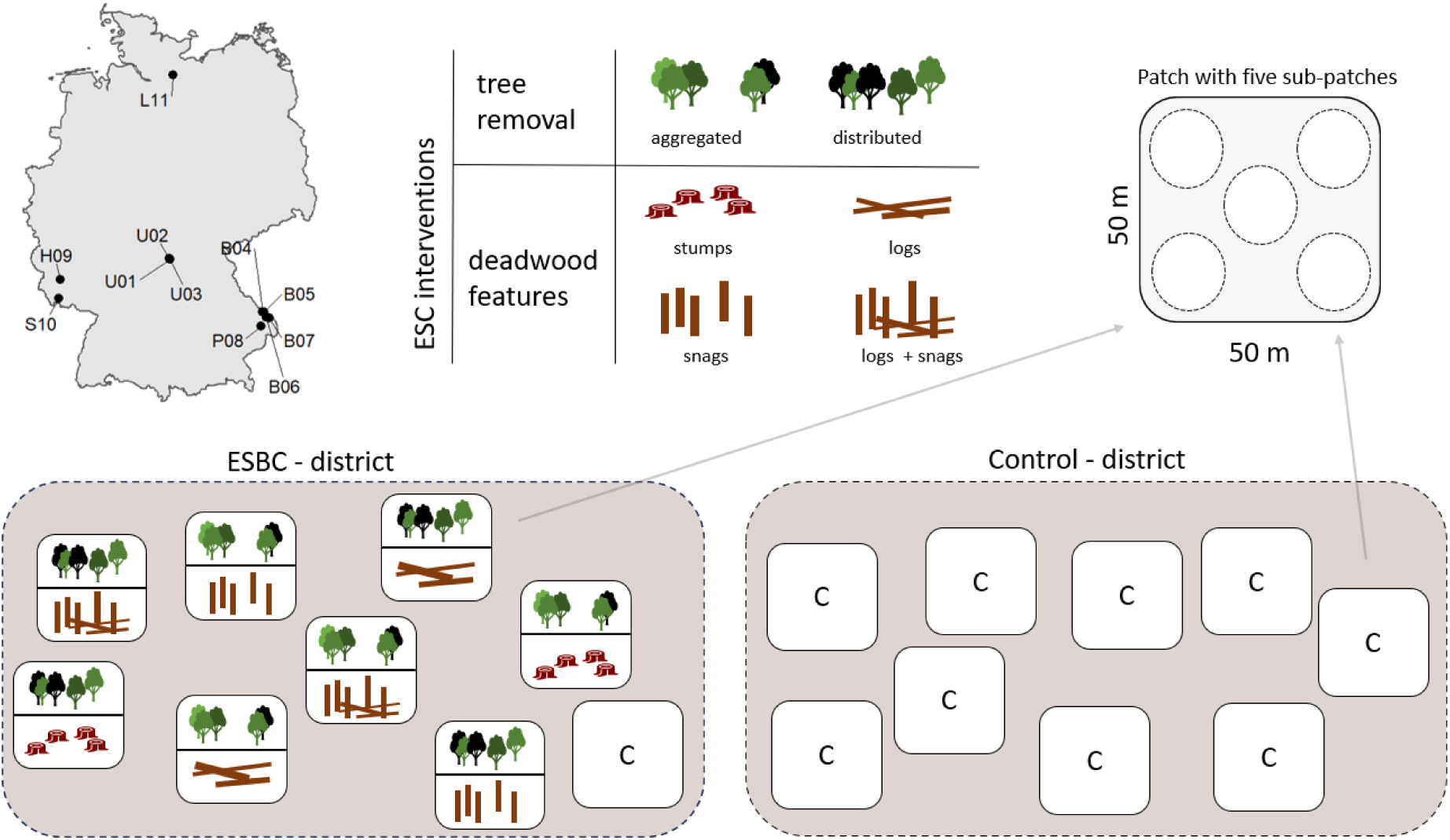
Conceptual figure showing the location of the 11 replicated experimental sites across Germany (map) and experimental design. Each experimental landscape consists of two paired forest districts: ESBC and control district. Districts comprised either nine patches (in eight of the 11 study sites) or 15 patches (in U01, U02, U03), with a patch size of 2.500 m² (50 m × 50 m) each. Within the ESBC districts, 8 (14 in U01, U02, U03) patches were treated experimentally (ESC) using a specific factorial combination of canopy structure (two levels: aggregated or distributed tree removal) and deadwood supply (four levels: stumps, logs, snags, or logs, and snags). Structural heterogeneity of the ESBC districts was further enhanced by adding one additional control patch per ESBC district.

### 2.3 Functional trait and phylogenetic data

To quantify functional diversity, we acquired four plant functional traits from Diáz et al. (2022). We selected traits related to the leaf economics spectrum and plant dispersal: leaf area, leaf nitrogen concentration, leaf mass per area, and diaspore mass. Plant height was excluded to avoid bias from measurements acquired on mature trees or shrubs that do not reflect the height of juveniles in the herb layer (up to 1 m). Missing trait values were imputed using a phylogenetically-informed approach with the funspace R package (Carmona et al. 2024).

For quantifying phylogenetic diversity, phylogenetic relatedness was assembled from established megatrees using the rtrees R package (Li, 2023).

### 2.4 Species’ forest affinities

Species were classified into four categories based on the strength of their association with forest habitats according to Heinken et al. (2022): Species primarily associated with closed-canopy forest environments (“closed forest specialists”; category 1.1), species primarily associated with forest edges and forest gaps (“open forest specialists”; category 1.2) and species that (primarily) occur in forests, but also in open habitats (“generalists”; categories 2.1 and 2.2). Species that are not associated with forest habitats were classified as “open habitat species” (category O). To select appropriate forest affinities, we used the categories “Germany, NW lowlands” for L11 and “Germany, uplands and mountains” for all other experimental landscapes. For 21 species, we used habitat associations listed in Flora Germanica (www.flora-germanica.de), as they were not included in the species list of Heinken et al. (2022). Across sites, 53.7% (123 species) of the species were classified as generalists, 31.9% (73 species) as closed forest specialists and 7.4% (17 species) as open forest specialists. Open habitat species accounted for 7% (16 species).

### 2.5 Light data

To explore how our treatments altered understorey light conditions and how these changes influenced understorey plant diversity, we took hemispheric photographs in each surveyed sub-patch (5 per patch) during summer 2023 using a camera with a fisheye lens (Nikon D7200, Sigma 4.5 mm F2.8 EX DC Circular Fisheye HSM lens). The camera was mounted on a tripod at a height of 1 m in the centre of each sub-patch and faced directly upwards. The diffuse site factor was calculated using programs developed by the chair of Photogrammetry (TUD Dresden University of Technology): “Segmentation” (v1.0.0.1) and “Hemisphere Tool” (v1.1 beta). The diffuse site factor represents the proportion of incident diffuse solar radiation that penetrates below the canopy, and thus corresponds to canopy openness. We used it as a proxy for understorey light availability in our study. The diffuse site factor is normalised to 1 (or 100%, respectively) across the entire image circle produced by a fisheye lens, which captures the full canopy above each point of interest.

### 2.6 Quantifying understorey plant diversity across spatial scales

#### Fundamental framework

We used a framework relying on Hill-Chao numbers to quantify changes in plant diversity with enhanced structural heterogeneity at multiple spatial scales. This framework allows us to differentiate between the responses of rare, common and dominant species to management interventions, which can provide important information about species assemblages. Hill numbers were originally developed based on species abundance/frequency data (Hill, 1973), but were adapted for incidence data, i.e. presence-absence data (Chao et al. 2014; Colwell et al. 2012). When using incidence data, Hill number of order q emphasizes species according to their detection frequency: infrequent species for q=0 (species richness), increasingly frequent species for q=1 (the exponential of Shannon entropy), and highly frequent species for q=2 (the inverse of Simpson concentration index).

Using incidence data can be necessary when the actual abundance cannot be reliably measured, as is often the case with plant species, where individual counts can be difficult to assess. Colwell et al. (2012) therefore recommend using incidence data, when independent sampling of individuals is not feasible. In our analyses of plant diversity, we converted cover values to incidence data. Species can thus have a value of either 1 (presence) or 0 (absence) in each of the 5 sub-plots per patch. Hill numbers were then computed from species incidence (detection) frequencies across sub-plots.

#### Three facets of diversity

A key advantage of Hill numbers is that they have been extended to a unified attribute diversity framework (Hill-Chao numbers; Chao et al. 2021), allowing quantification of three major facets of diversity. Under the extended framework, taxonomic diversity (TD) quantifies the effective number of equally-abundant species; (mean) phylogenetic diversity (PD) quantifies the effective number of equally-divergent lineages when the phylogenetic tree depth is scaled to 1; functional diversity (FD) quantifies the effective number of equally distinct virtual functional groups (or functional “species”) based on species’ functional traits. FD standardisation requires a sufficient number of sampling units (subplots, in our cases) to obtain reliable results (Colwell and Chao, 2022). Because our dataset included only 5 subplots, we treated species detection frequency as a proxy for species abundance and obtained approximate FD standardized estimates. Because all three facets are expressed in the same unit of species or lineage equivalents, they are directly comparable (Chao et al. 2021).

#### Multiple spatial scales (alpha, gamma, and beta diversity)

In our hierarchical diversity analysis, alpha diversity quantifies mean diversity at the patch level and can be interpreted as the effective number of species/lineages/functional groups per patch (Chiu et al. 2014; Chao et al. 2023). Gamma diversity quantifies diversity at the district level (control or ESBC, each consisting of 9 or 15 patches) and represents the effective number of species/lineages/functional groups in a district. Beta diversity quantifies compositional differentiation among patches, following Whittaker’s (1960) multiplicative framework but generalized via Hill–Chao numbers for any order *q*. Beta diversity can be interpreted as the effective number of non-overlapping patches (patch equivalents) and is bounded between 1 (all patches identical) and the total number of patches (9 or 15; no shared species).

To account for differences in patch number across sites, we applied the 1-S-transformation (Table 1 of Chao et al. 2019) to convert beta diversity into a Jaccard-type dissimilarity index. This turnover measure (*sensu* Lande 1996, i.e. turnover = gamma - alpha) is expressed relative to gamma diversity and standardised by the number of patches. It ranges from 0 (identical patches) and 1 (no shared species among patches), enabling direct comparisons across sites.

### 2.7 Statistical standardisation and meta analyses

Observed diversity based on sampling data are sensitive to both sampling effort and sample completeness, making meaningful comparisons among sites challenging. Therefore, standardising diversity estimates for sample coverage, which is a measure of sample completeness defined as the proportion of overall incidences (occurrences) that belong to observed species (Chiu 2023), is essential for obtaining reliable diversity estimates (Kortmann et al. 2025). The concept of sample coverage was originally developed by Alan Turing in his WWII code-breaking work, and has since been adopted by ecologists as a basis for standardizing samples (Chao et al. 2014).

All 22 forest districts exhibited incomplete data (sample coverage < 1; Figure S1, S2) with observed coverage ranging from 0.61 to 0.90. To account for the dependency of observed diversity on sample coverage, we standardised all 22 districts to a fixed level of sample coverage of 0.9 using rarefaction and extrapolation (Figure S3), following the procedures in iNEXT.beta3D (Chao et al. 2023). Standardization was performed for all three diversity facets and for the alpha, gamma, and beta components.

Based on the coverage-standardised estimates, we applied a meta-analysis approach to evaluate how enhanced structural heterogeneity affected gamma diversity across the three diversity facets. To further identify drivers of change in gamma diversity, we extended the meta-analysis to the corresponding alpha and beta components. This meta-analytical framework including coverage-based standardization, meta-analysis, and comparison across diversity orders (q=0, q=1, q=2) and facets represents a novel approach for cross-site comparisons under differing environmental contexts and levels of sample completeness.

For each of the 11 sites, the ESBC effect was estimated as the difference in standardised gamma/alpha/beta diversity between ESBC and control districts for each diversity facet and each Hill-Chao number (q=0, q=1, q=2). Uncertainty was assessed using a bootstrap procedure with 50 replications to generate 95% confidence intervals (see Chao et al. 2014 for details). For any specified spatial scale, diversity facet, and value of q, the ESBC-effect values across sites were then weighed by the inverse of their variances (i.e., the higher the variance, the lower the weight) to obtain the overall ESBC effect and its confidence interval. The overall effect was considered significant when the 95% confidence interval did not include zero. All computation and graphics were performed using the new function, iNEXTmeta_beta, adapted from iNEXT.beta3D (Chao et al. 2023). The function is currently available at: https://github.com/AnneChao/iNEXT.meta.

### 2.8 Further data analyses

To assess how species with different forest affinities responded to the ESBC treatment, we calculated for each forest affinity category the difference in the total number of observed species between each pair of control and ESBC district. Confidence intervals were derived using a bootstrapping approach included in the *Hmisc* package in R (Harrel 2025).

We used structural equation modelling (SEM) to evaluate the drivers of plant diversity at the landscape scale. Specifically, we aimed to disentangle the direct effects of increased structural heterogeneity on taxonomic plant diversity at multiple spatial scales from indirect effects mediated by changes in light availability and heterogeneity. For each district, light availability was quantified by first calculating the mean of five diffuse site factor measurements per patch, and then computing the mean across all patches within the district. Light heterogeneity was quantified as the coefficient of variation (CV) of the diffuse site factor based on five sub-patch measurements per patch, and averaged across all patches within a district resulting in one single value per district. The effect of treatment (binary variable) was coded as 0 = control and 1 = ESBC and then treated as a numeric covariate (Rosseel 2012). Thus positive relationships indicate a positive effect of ESBC on light conditions (i.e., light availability and heterogeneity) and plant diversity. A significant direct effect of treatment on regional plant diversity would suggest that gamma diversity is primarily driven by differences among communities (i.e., beta diversity). In contrast, a significant indirect effect of treatment mediated through mean local plant diversity (alpha diversity) would indicate that changes in gamma diversity are largely attributable to shifts in average alpha diversity within a district. We used standardised diversity values for taxonomic diversity of the order q=0 obtained from the meta-analysis. Alternative SEMs using higher orders of q (q=1 and q=2) gave qualitatively similar results (Figures S3-S4). The SEMs were fitted using the ‘piecewiseSEM’ framework (Lefcheck, 2016), based on linear models. Note that the standardised diversity values represent continuous values and are not restricted to integers. Path coefficients were standardised via z-transformation to facilitate a direct comparison of the relative strengths of individual pathways. Model fit was assessed using Fisher’s *C* statistics, with p-values > 0.05 indicating an adequate fit to the data (Lefcheck, 2016).

All analyses were performed using R (version 4.3.2, R Core team 2025) and figures were created using *ggplot2* (Wickham 2016).

## 3. Results

### 3.1 Consistent increases in gamma and alpha, but not beta diversity in structurally heterogeneous forests

Overall, the meta-analysis revealed that the enhancement of structural heterogeneity significantly increased the taxonomic diversity of understorey plants at the gamma and alpha scales across all experimental landscapes (Figure 2). Compared to control districts, ESBC districts contained on average 20 more effective species (95% CI: 16.7-24.0), including 14 (95% CI: 13.2-15.8) that were highly frequent. At q=0, the effect was consistently positive across all 11 experimental landscapes and statistically significant in all but three cases (Figure S5). A similar pattern was evident at the alpha scale: patches within ESBC districts contained on average 8 additional effective species (95% CI: 7.3–9.0), including 5 (95% CI: 4.8–5.4) that were highly frequent. This positive effect of ESBC on local plant species richness (q=0) was detected in all sites and reached significance in 8 of them (Figure S6).

**Figure 2:**
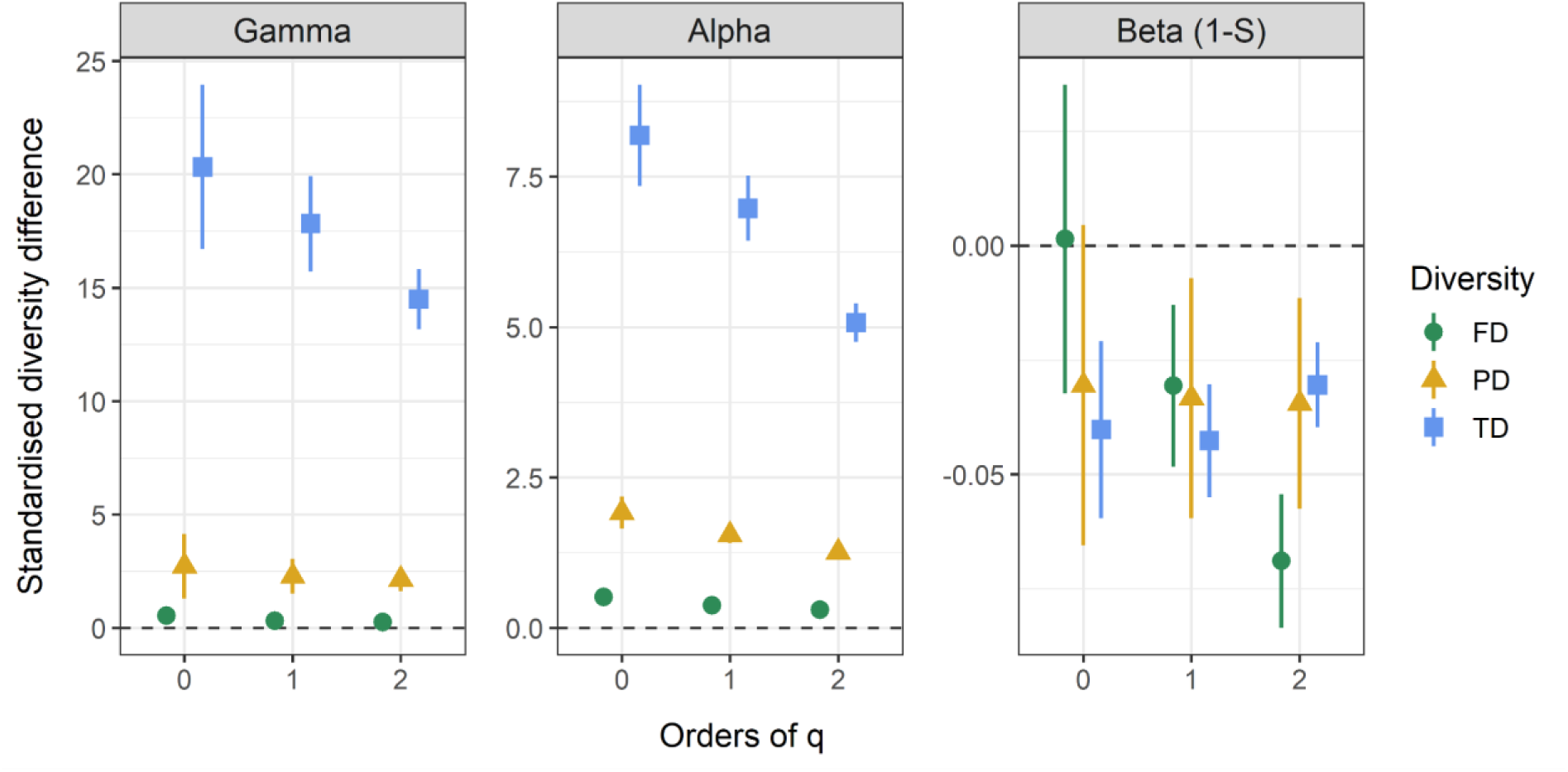
Meta-analysis results across 11 experimental landscapes. Symbols indicate the mean predicted changes in standardised diversity between heterogeneous (ESBC) and homogenous (control) forest districts. Positive values indicate higher diversity in ESBC compared to control districts, with effects considered significant when the 95% confidence intervals (error bars) do not include zero. The standardised difference in plant diversity was calculated between each pair of control and ESBC district in directly comparable species-equivalent units for gamma and alpha diversity and for different diversity facets (taxonomic, phylogenetic and functional diversity) and diversity orders (q=0, q=1 and q=2). A Jaccard-type turnover transformation (1-S) of multiplicative beta diversity was applied to correct for differences in the number of assemblages (i.e. 9 or 15) sampled in the different experimental landscapes. See forest plots in supplement (Figures S5 - S31) for detailed results.

The increase in taxonomic diversity observed in ESBC districts was accompanied by an increase in phylogenetic and functional diversity at both the gamma and alpha scales (Figure 2). At the gamma scale (forest district), standardised differences in phylogenetic and functional diversity were consistent across diversity orders, indicating that the gains were mainly driven by highly frequent lineages and functional groups. At the alpha scale (forest patch), however, standardised differences varied more strongly with diversity order, suggesting that the increase reflected contributions from both highly frequent and less frequent lineages and functional groups.

In contrast to gamma and alpha diversity, an enhancement of structural heterogeneity resulted in a slight decrease of beta diversity on average for all facets and diversity orders, except for functional and phylogenetic diversity of order q=0 (Figure 2). Across sites and diversity orders, we observed a strong variation in the effect of the ESBC treatment on the taxonomic (Figures S7, S10, S13), phylogenetic (Figures S16, S19, S22), and functional (Figures S28 and S31) beta diversity of understorey plants. For example, while taxonomic beta diversity (q=0) significantly increased in four sites (B05, B06, H09, U03), we also observed a significant decrease in beta diversity in four sites (B04, S10, U01, U02) and no change in beta diversity in three sites (B07, L11, P08) (Figure S7).

Species gains for all diversity facets were unevenly distributed along diversity orders for alpha and gamma diversity, but differences between diversity orders were most pronounced for taxonomic diversity. For gamma and alpha diversity of all diversity facets, infrequent species (q=0) showed the strongest response to enhanced heterogeneity followed by more frequently occurring species (q=1) and highly frequent species (q=2). For beta diversity, mean effect sizes were negative across diversity facets and orders with the exception of phylogenetic and functional diversity of order q=0, where no significant effects were detected. For functional beta diversity, mean effect sizes were neutral for q=0 (0.001) or slightly negative for q=2 (-0.07). For phylogenetic beta diversity, mean differences were very similar between diversity orders, but showed a slight decrease (increase in effect sites), from q=0 to q=2 (-0.031 for q=0, -0.034 for q=2). For taxonomic diversity, the highest average difference between control and ESBC district was observed for q=1 (-0.033). Notably, while most of the different experimental landscapes showed clear increases in gamma and alpha diversity, there was no consistent pattern across landscapes for beta diversity (Figures S5 - S31).

Absolute standardized gamma diversity and alpha diversity differed between experimental landscapes and control districts for the different diversity orders (Figures S32, S33), but the magnitude of the treatment effect on gamma and alpha diversity was consistently highest for taxonomic diversity, followed by phylogenetic and functional diversity, corroborating a robust pattern across the different diversity facets. The observed increases in alpha diversity, coupled with slight decreases or neutral responses for PD and FD at q=0 in beta diversity across diversity facets and diversity orders, indicate that gamma diversity was primarily driven by changes in alpha diversity.

### 3.2 Gamma diversity in ESBC districts driven by alpha diversity via landscape light heterogeneity

We found that the observed increase in gamma diversity in the ESBC districts was primarily driven by higher alpha diversity on average (standardised path coefficient = 0.87, p <0.001), rather than by increased species turnover between patches (i.e., beta diversity), as indicated by the non-significant direct effect of ESBC treatment on gamma diversity (standardised path coefficient = 0.07, p = 0.500; Figure 4). This effect was largely mediated by greater between-patch light heterogeneity in ESBC districts (landscape light heterogeneity; standardised path coefficient = 0.51, p = 0.042), but not by their mean light availability (landscape light availability; standardised path coefficient = 017, p = 0.515). Moreover, changes in landscape light availability and heterogeneity were not directly related to changes in gamma diversity (standardised path coefficients = -0.02 and 0.10, p = 0.815 and 0.349). This suggests that the observed positive effects of increased structural heterogeneity on gamma diversity are primarily driven by local increases in species diversity, likely due to greater variety of light niches in structurally heterogeneous forest landscapes.

**Figure 3:**
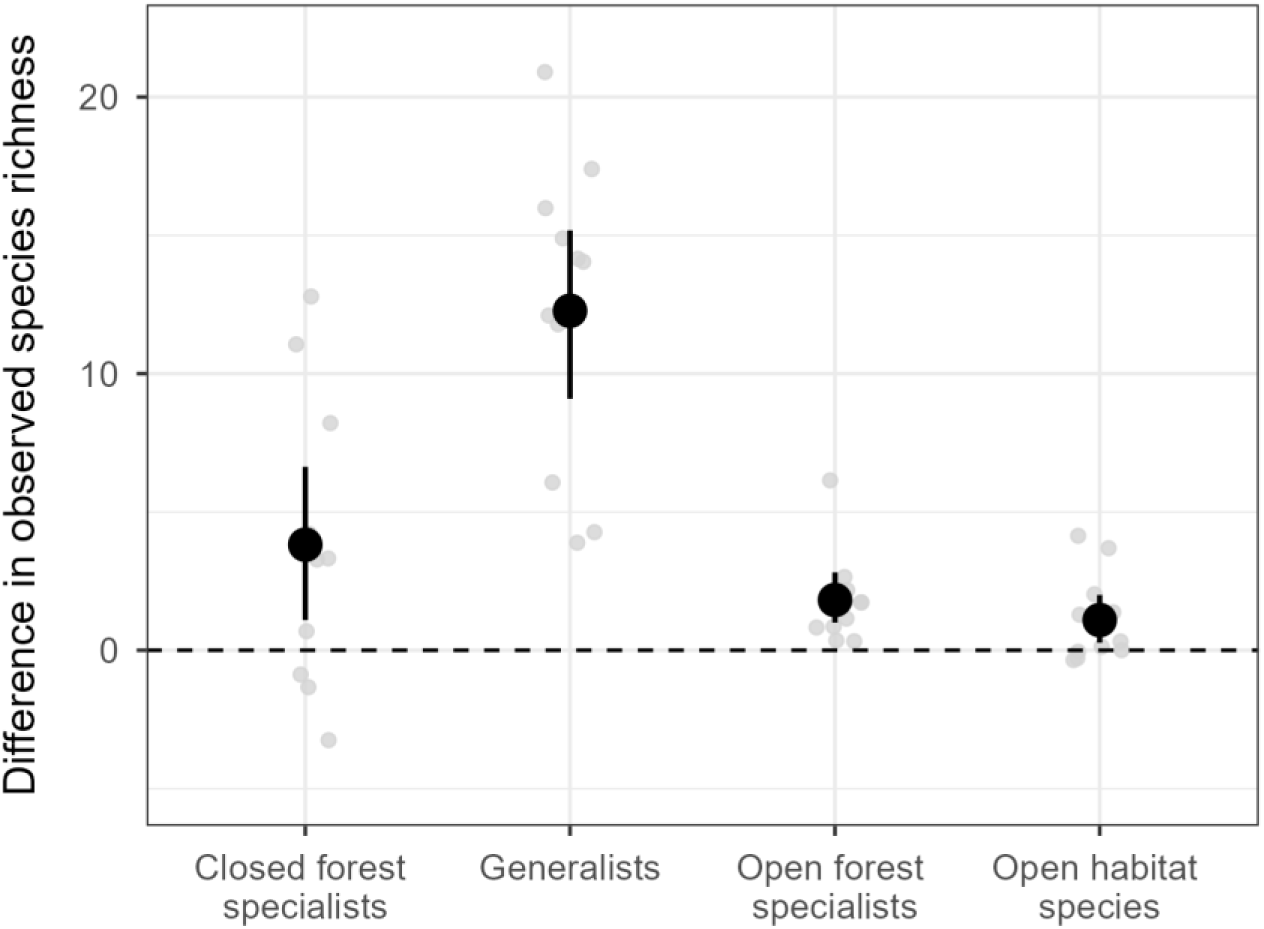
Effects of an enhancement of structural heterogeneity in forest landscapes on species’ affinity to forests. Light grey points represent the differences in the observed species richness per forest affinity category between the ESBC and control district across experimental landscapes, and error bars denote the 95%-confidence intervals computed by bootstrapping. Species are categorised as closed forest specialists, open forest specialists, generalists and open habitat species. Positive differences denote higher observed species richness in ESBC districts, while negative differences denote the opposite. Note, that a difference of zero (in this dataset) indicates that the category was absent in both districts.

**Figure 4:**
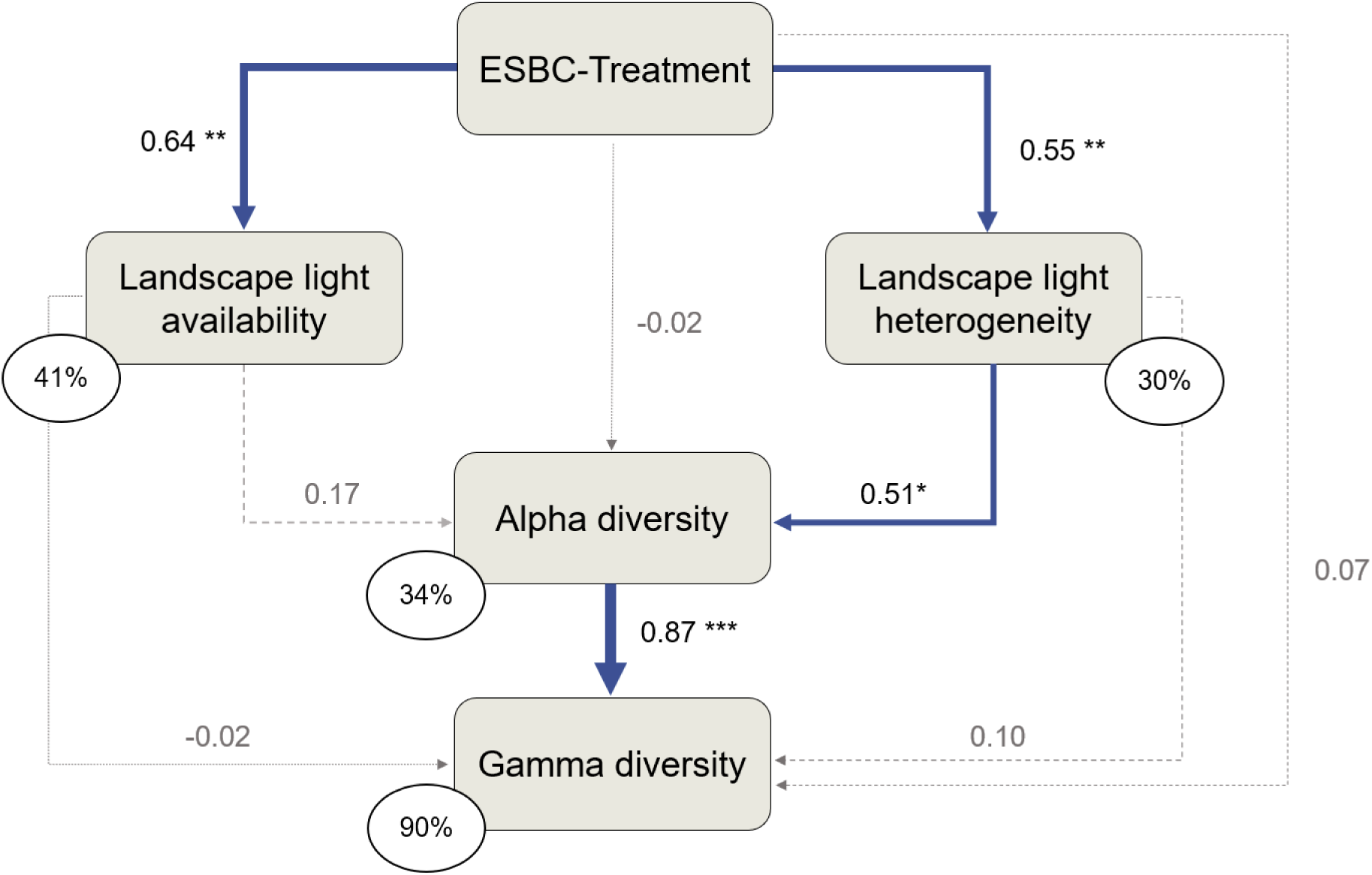
Structural equation model linking understorey plant diversity and management-induced changes in light conditions across multiple spatial scales. The models were fitted using standardised taxonomic diversity values of the diversity order q=0 for alpha diversity and gamma diversity. Positive relationships between enhanced structural heterogeneity between forest patches (ESBC) and light conditions (light availability and heterogeneity) or plant diversity (mean joint alpha and gamma diversity) indicate a positive ESBC effect. Landscape light availability refers to the mean canopy openness of all patches within a district (mean within-patch canopy openness), while landscape light heterogeneity refers to the coefficient of variation in canopy openness across patches within a district (between-patch canopy openness heterogeneity). Blue arrows indicate significant (* p < 0.05, ** p < 0.01 and *** p < 0.001) and dashed grey arrows non-significant (p > 0.05) relationships. Numbers next to arrows are standardised path coefficients and arrow width is proportional to their effect size. Percentage values are explained variances of endogenous variables. The model provided a good fit to the data: Fisher’s C = 1.072, df = 2, p-value = 0.585.

### 3.3 Positive effects of ESBC treatment on both forest generalists and specialists

On average, enhancing structural heterogeneity at the landscape scale (ESBC) led to increases in closed and open forest specialists, generalists, and open habitat species (Figure 3). The highest absolute gains were observed for generalist species (mean = 12.3, CI: 9.4 - 15.3), which were significantly greater than increases in the remaining categories (+322% to +554%). Absolute gains in closed forest specialists (mean = 3.8, CI: 1.2 - 6.7) were approximately two-fold higher than gains in open forest specialists (mean = 1.8, CI: 1.0 - 2.8) and open habitat species (mean = 1.1, CI: 0.3 - 2.1).

## 4. Discussion

Our findings provide experimental evidence that management promoting structural heterogeneity in forest landscapes can play a fundamental role in maintaining understorey plant diversity. We show that an enhancement of structural heterogeneity between forest patches increases the taxonomic, phylogenetic, and functional diversity of understorey plants. Since plant diversity increased not only among generalist or open habitat species but also among forest specialists, these results highlight the potential of structural interventions as an effective conservation tool in managed temperate forests.

### 4.1 Structural heterogeneity between forest patches enhances gamma diversity through increased local diversity

Experimentally assessing heterogeneity–diversity relationships at different spatial scales is highly relevant for understanding biodiversity patterns and underlying ecological processes at larger scales (Bartels & Chen 2010; Heidrich et al. 2020; Stein et al. 2014). If habitat conditions change, plant community composition can show rapid responses with studies reporting effects on species richness (Heidrich et al. 2020) but also shifts in abundances (Dornelas et al. 2009). Our results reveal significant effects on both richness and frequency (as a proxy for abundance) across the alpha and gamma scales. As hypothesised (H1), gamma diversity of understorey vegetation was higher in structurally heterogeneous landscapes (ESBC districts) than in homogenous ones across a wide range of environmental conditions. Our experimental evidence coincides with findings of a modelling study, which suggested that the heterogeneity of different forest management systems within a ‘virtual’ (i.e. modelled) forest landscape across Germany strongly promotes regional vascular plant diversity (gamma diversity) due to a higher spatial heterogeneity in habitat conditions among stands (Schall et al. 2020). Moreover, we found that the higher gamma diversity in ESBC districts was primarily driven by an overall increase in alpha diversity, indicating that local structural interventions can scale up to promote species gains at larger spatial scales. Importantly, this effect was positively associated with variation in light availability between forest patches. While our two contrasting tree-removal treatments (aggregated vs. distributed) were applied at the local scale and mainly increased light heterogeneity within individual patches, combining all patches generated a broader gradient of light conditions within an ESBC district. As a result, both shade-adapted species and those favouring higher light exposure could find suitable habitats, likely due to the greater variety of light niches, providing empirical support for the habitat heterogeneity hypothesis. Notably, we found that light heterogeneity at the landscape scale, rather than mean light availability, was the main driver for changes in overall alpha diversity. In contrast, Dormann et al. (2020) found that local plant species richness in temperate forest understoreys increased with light availability, but not with light heterogeneity, suggesting that the effects of light heterogeneity on diversity patterns may emerge primarily at larger spatial scales.

There are several pathways through which new species can enter patches. One could be through germination of seeds that were already present in the soil seedbank and found suitable conditions under altered canopy structures - conditions that would naturally occur following disturbances (Plue et al. 2021). The observed increase in gamma diversity further suggests that some species are either newly established within the landscape or have colonised patches from the surrounding matrix. This aligns with the metacommunity framework (Leibold and Chase 2017), which suggests that species can recruit and persist in less suitable habitats through continuous immigration (i.e., mass effects). Thus, local populations may benefit from such source-sink dynamics within heterogeneous landscapes, which could explain the overall increase in alpha diversity. These new species may occur in some patches or across multiple patches, as indicated by the high gains in highly frequent species, thus contributing to an increase in gamma diversity, while also potentially contributing to a decrease in beta diversity. However, many forest species are known to be slow dispersers and display rather conservative traits (Blondeel et al. 2020; Brunet and von Oheimb 1998, de Pauw et al. 2021). Thus, responses of especially slow dispersers may be lagging which may suggest that effects on the already initially high beta diversity take longer to manifest. Additionally, many forest specialists are characterised by low seed production and have only short time frames for dispersal to maintain the ability to germinate, while generalist species often produce a large amount of long-lived seeds and have strong competitive dispersal abilities, likely leading to higher representation of generalist species in the soil seed bank (Büchi and Vuilleumier 2014, Whigham 2004).

### 4.2 Negative overall effect on beta diversity, but lack of generality across experimental landscapes

On average, beta diversity showed a weak negative response across diversity facets and orders indicating slight homogenisation rather than differentiation, in contrast to our second hypothesis (H2). Similarly, Cramer and Willig (2005) observed homogenisation in small mammals, which they attributed to edge effects at the interface of contrasting habitats, known for supporting higher diversity. An increase in habitat diversity within patches may explain the slight homogenisation between patches, as species with different habitat requirements can coexist in structurally heterogeneous patches, which is in line with the observed increase in species richness across forest affinity categories. However, it is important to note that the small effect size of beta diversity should be taken into account when assessing the magnitude of homogenisation and its ecological relevance. Importantly, the low effect sizes in differences should not be mistaken as an overall low beta diversity. On the contrary, taxonomic beta-diversity of order q=0 ranges from 0.525 to 0.858 (weighted mean = 0.659) in control districts and 0.222 to 0.866 (weighted mean = 0.619) in ESBC districts (Figure S34). This indicates an overall high beta diversity regardless of the treatment, decreasing only slightly but remaining high in structurally enhanced districts. A possible explanation for this weak effect can be attributed to the fact that our treatments involved two canopy structure modifications, but four to six manipulations of deadwood structures. We showed that heterogeneous light conditions had a strong impact on understorey diversity patterns in the ESBC districts, consistent with the widely recognised importance of light in determining understorey plant diversity and community composition (Helbach et al. 2022; Dormann et al. 2020). We therefore assume that the effect of deadwood, as a second manipulated aspect of forest structure, was likely neglectable in our study, especially at this rather young decomposition stage (Sandström et al. 2019).

Our results also demonstrate that both the magnitude and direction of beta diversity responses to enhanced structural heterogeneity can vary considerably across large-scale gradients of environmental heterogeneity (i.e., across landscapes). This implies that single case studies could come to overly simplified conclusions that are specific to the environmental context of the study. There is growing recognition for a lack of generality in beta diversity patterns as highlighted in two recent meta-analyses (Keck et al. 2025; Gonçalves-Souza et al. 2025). Here, we experimentally show that this pattern persists even when correcting for sample coverage and analysing across diversity facets. Different responses of individual district-pairs can be the result of many factors. Sola and Griffin (2025) concluded that the influence of (local) heterogeneity on biodiversity depends on how it is generated and measured, the taxa considered, the organisms’ traits and prevailing environmental conditions. In our study, experimental landscapes exhibiting comparatively high diversity in the control district tend to gain fewer species than those with lower reference diversity (Figure S35). On the other hand, increases in few species have a comparably greater impact on multiplicative beta diversity and its 1-S-transformation in species-poor landscapes. Other factors shaping patterns in individual experimental landscapes may include edaphic conditions, dispersal limits, or land-use history (Borderieux et al. 2023). Site-specific contexts, such as the regional species pool or initial heterogeneity, can also influence outcomes. Still, consistent alpha and gamma diversity trends suggest general mechanisms by which increased structural heterogeneity supports understorey plant diversity across landscapes.

### 4.3 Greatest increases for taxonomic diversity suggest redundancy in increases of functional and phylogenetic diversity

We consistently observed the greatest increases in taxonomic diversity, followed by phylogenetic and functional diversity. This pattern suggests that enhancing structural heterogeneity between forest patches primarily benefits taxonomic diversity, while having smaller effects on trait and lineage diversity. On the other hand, taxonomic diversity itself is much higher than phylogenetic and functional diversity. In effective species numbers, 103.89 species in the ESBC district of U03 (q=0), for example, relate to 16.35 lineages and 3.65 functional groups, suggesting a high redundancy in traits and a high degree of shared evolutionary history between species (Figures S5, S14, S23). The overall lower diversity in functional groups and lineages compared to taxonomic diversity (Figure S32) suggests a rather strong selection effect on specific lineages/traits, leading to high functional and phylogenetic similarity between species and explaining the low effect sizes for functional and phylogenetic diversity.

Padullés Cubino et al. (2023) found that species loss due to global change in temperate forest understories may be related to a loss of specific phylogenetic branches and specific ecological strategies. While our method does not allow for the identification of such branches, we were able to show that changes in phylogenetic diversity differ from those in taxonomic diversity. Our results showed smaller effect sizes for phylogenetic diversity than for taxonomic diversity. However, this low increase might be attributable to the generally limited number of phylogenetic lineages present as calculated in our framework. Nonetheless, this indicates that we likely gained species within existing branches, rather than species from distinct lineages.

When comparing patterns in effect sizes (q=0 > q=1 > q=2) between diversity facets, beta diversity showed the greatest variation across diversity orders. Notably, only functional diversity showed a pattern of increasingly negative differences in beta diversity with higher diversity order, meaning that differences in functional diversity are most pronounced for more frequently occurring species. However, we also observe negative differences for functional diversity for some experimental landscapes at q=0 and q=1. As stated before, functional diversity needs to be interpreted with care due to sparse data.

### 4.4 Increase in species richness across forest affinity categories

Habitat changes are linked to shifts in community composition that often benefit certain species at the cost of others through environmental filtering and niche processes (Keck et al. 2025). The observed increase in alpha and gamma diversity in structurally enhanced forest districts was driven by increases in observed species richness across all categories of forest affinity. We observed the strongest absolute increase among generalist species, as those species were most frequent across experimental landscapes due to their less specific habitat requirements, which enabled them to recruit and exploit a wider range of habitats and resources (Stein et al. 2014; de Pauw et al. 2021). In line with hypothesis 3 (H3), this pattern also holds when looking at relative differences between ESBC and control districts, but differences were less distinct (Figure S36). Moreover, in terms of absolute differences, we found that both forest specialists of closed and open forest habitats benefitted from an enhanced structural heterogeneity in ESBC districts with effects being stronger for closed forest specialists. For example, it is conceivable that contrasting environmental conditions (e.g., the spatial variation of high-light and low-light patches) can foster niche partitioning, thereby reducing competitive pressure between forest specialists and generalists (Lorer et al. 2024).

Categories of forest affinity are closely linked to preferences in light conditions as light availability is a strong filter for understorey plant communities (Heinken et al. 2022). European broadleaved forests are naturally subject to manifold disturbances, resulting in a mosaic of several successional stages nowadays often suppressed by forest management (Hilmers et al. 2018; Senf and Seidl 2021). More than 80% of European temperate forest plants, however, are associated with high light availability and show a preference for heterogeneous light conditions in forests, challenging the prevailing closed-forest paradigm (Czyzewski and Svenning, 2025). This is mirrored in the forest affinity list by Heinken et al. (2022), where proportionally, on the European level, generalist species outnumber specialist forest species by far, explaining the observed high numbers of generalist species in our data. While generalist species are often fast colonisers, forest specialists are known to be rather slow colonisers (de Pauw et al. 2021). Given that our treatments mimic diverse successional stages within a landscape, we observe stronger increases of species richness of all categories of forest affinity in structurally diverse forests. Notably, we do not observe a disproportionate gain of species with more open habitat preferences (e.g. grassland associated species), indicating that our management interventions do not shift plant community composition in favor of open habitat species.

### 4.5 Implications for forest conservation and restoration

Our results demonstrate that the variation of small-scale interventions enhancing structural heterogeneity between forest patches strongly favours understorey plant diversity in forest landscapes across diversity facets. These gains included both forest specialists and generalist species. Moreover, we found that the increase in taxonomic, functional, and phylogenetic gamma diversity was strongly associated with more heterogeneous light conditions within a landscape. Consequently, landscape mosaics of canopy-manipulating interventions, such as combined single tree removal and group felling, allow forest management to mimic early and late forest development stages associated with open forest habitats (Vild et al. 2024), which support the highest levels of plant diversity typical of forest ecosystems (Hilmers et al. 2018). These findings demonstrate that management can enhance understorey plant diversity by increasing structural heterogeneity, providing a complementary strategy that supports, but does not replace, the vital role of unmanaged forests in conserving forest biodiversity.

## Acknowledgements

We want to acknowledge the support of several student research assistants and technicians during vegetation surveys and field campaigns. We further thank the entire team of the BETA-FOR research unit for their support. This work was funded by the German Research Foundation (DFG-FOR 5375; project nr. 459717468).

