## Supplement for "Enhancing structural heterogeneity in managed forest landscapes promotes gamma but not beta diversity in understorey plant communities"

Bradler et al.

**Table S1:** Overview of study sites with elevation above sea level (NN) in m: mean, minimum and maximum elevation above sea level. Treatment-establishment (ESBC): 2015/2016: B04, B05, B06, B07, P08, S10; 2017/2017: L11; 2018/2019: U01, U02, U03.

| Site | District | Mean NN | Min NN | Max NN | Vegetation Type |
|---|---|---|---|---|---|
| B04 | Control | 941 | 892 | 987 | Luzulo-Fagetum |
| B04 | ESBC | 835 | 796 | 871 | Luzulo-Fagetum |
| B05 | Control | 1114 | 1086 | 1144 | Luzulo-Fagetum |
| B05 | ESBC | 1046 | 1004 | 1077 | Luzulo-Fagetum |
| B06 | Control | 809 | 780 | 836 | Luzulo-Fagetum |
| B06 | ESBC | 850 | 815 | 873 | Luzulo-Fagetum |
| B07 | Control | 971 | 935 | 992 | Luzulo-Fagetum |
| B07 | ESBC | 885 | 836 | 917 | Luzulo-Fagetum |
| H09 | Control | 652 | 642 | 662 | Galio-Fagetum |
| H09 | ESBC | 656 | 652 | 666 | Galio-Fagetum |
| L11 | Control | 43 | 42 | 43 | Galio-Fagetum |
| L11 | ESBC | 40 | 38 | 42 | Galio-Fagetum |
| P08 | Control | 508 | 450 | 532 | Luzulo-Fagetum |
| P08 | ESBC | 484 | 453 | 526 | Luzulo-Fagetum |
| S10 | Control | 287 | 260 | 296 | Galio-Fagetum |
| S10 | ESBC | 304 | 278 | 318 | Galio-Fagetum |
| U01 | Control | 375 | 362 | 393 | Hordelymo-Fagetum |
| U01 | ESBC | 322 | 296 | 342 | Hordelymo-Fagetum |
| U02 | Control | 363 | 350 | 374 | Hordelymo-Fagetum |
| U02 | ESBC | 335 | 299 | 362 | Hordelymo-Fagetum |
| U03 | Control | 309 | 299 | 330 | Hordelymo-Fagetum |
| U03 | ESBC | 313 | 300 | 330 | Hordelymo-Fagetum |

**Table S2:** Overview of tree species composition per site and district. Values are average basal area (%) across districts (Control or ESBC).

| Site | District | Species | Cover |
|---|---|---|---|
| B04 | Control | Acer sp. | 8.72 |
| B04 | Control | Fagus sylvatica | 83.93 |
| B04 | Control | Picea abies | 6.55 |
| B04 | Control | Pseudotsuga menziesii | 0.8 |
| B04 | ESBC | Acer sp. | 0.34 |
| B04 | ESBC | Betula pendula | 0.19 |
| B04 | ESBC | Fagus sylvatica | 91.97 |
| B04 | ESBC | Fraxinus excelsior | 0.18 |
| B04 | ESBC | Picea abies | 7.18 |
| B04 | ESBC | Sorbus aucuparia | 0.14 |
| B05 | Control | Acer sp. | 0.07 |
| B05 | Control | Fagus sylvatica | 91.65 |
| B05 | Control | Picea abies | 7.8 |
| B05 | Control | Sorbus aucuparia | 0.21 |
| B05 | Control | Tilia sp. | 0.27 |
| B05 | ESBC | Acer sp. | 1.01 |
| B05 | ESBC | Fagus sylvatica | 81.53 |
| B05 | ESBC | Picea abies | 17.2 |
| B05 | ESBC | Sorbus aucuparia | 0.26 |
| B06 | Control | Acer sp. | 3.43 |
| B06 | Control | Betula pendula | 0.02 |
| B06 | Control | Fagus sylvatica | 88.17 |
| B06 | Control | Picea abies | 8.09 |
| B06 | Control | Populus tremula | 0.24 |
| B06 | Control | Prunus avium | 0.05 |
| B06 | ESBC | Abies alba | 3.67 |
| B06 | ESBC | Acer sp. | 0.19 |
| B06 | ESBC | Betula pendula | 0.11 |
| B06 | ESBC | Fagus sylvatica | 83.39 |
| B06 | ESBC | Picea abies | 12.43 |
| B06 | ESBC | Sorbus aucuparia | 0.21 |
| B07 | Control | Acer sp. | 0.35 |
| B07 | Control | Fagus sylvatica | 66.89 |
| B07 | Control | Picea abies | 32.77 |
| B07 | ESBC | Acer sp. | 1.34 |
| B07 | ESBC | Fagus sylvatica | 68.56 |
| B07 | ESBC | Picea abies | 29.68 |
| B07 | ESBC | Sorbus aucuparia | 0.41 |
| H09 | Control | Carpinus betulus | 0.05 |
| H09 | Control | Fagus sylvatica | 98.03 |
| H09 | Control | Picea abies | 1.93 |
| H09 | ESBC | Fagus sylvatica | 99.46 |
| H09 | ESBC | Picea abies | 0.54 |
| L11 | Control | Acer sp. | 0.79 |
| L11 | Control | Carpinus betulus | 0.04 |

| | | | |
|---|---|---|---|
| L11 | Control | Fagus sylvatica | 88.75 |
| L11 | Control | Larix decidua | 0.29 |
| L11 | Control | Quercus sp. | 10.12 |
| L11 | ESBC | Acer sp. | 0.2 |
| L11 | ESBC | Carpinus betulus | 1.55 |
| L11 | ESBC | Fagus sylvatica | 80.41 |
| L11 | ESBC | Larix decidua | 1.83 |
| L11 | ESBC | Picea abies | 1.08 |
| L11 | ESBC | Quercus sp. | 14.92 |
| P08 | Control | Betula pendula | 0.77 |
| P08 | Control | Fagus sylvatica | 71.9 |
| P08 | Control | Picea abies | 21.02 |
| P08 | Control | Pinus sylvestris | 6.31 |
| P08 | ESBC | Abies alba | 0.27 |
| P08 | ESBC | Fagus sylvatica | 78.28 |
| P08 | ESBC | Picea abies | 19.7 |
| P08 | ESBC | Pinus sylvestris | 1.63 |
| P08 | ESBC | Prunus avium | 0.06 |
| P08 | ESBC | Quercus sp. | 0.06 |
| P08 | ESBC | Sorbus aucuparia | 0.01 |
| S10 | Control | Acer sp. | 0.35 |
| S10 | Control | Betula pendula | 1.19 |
| S10 | Control | Carpinus betulus | 15.89 |
| S10 | Control | Fagus sylvatica | 40.09 |
| S10 | Control | Larix decidua | 2.69 |
| S10 | Control | Picea abies | 5.8 |
| S10 | Control | Pinus sylvestris | 0.52 |
| S10 | Control | Populus tremula | 0.09 |
| S10 | Control | Prunus avium | 0.19 |
| S10 | Control | Pseudotsuga menziesii | 0.01 |
| S10 | Control | Quercus sp. | 33.09 |
| S10 | Control | Salix sp. | 0.03 |
| S10 | Control | Ulmus sp. | 0.06 |
| S10 | ESBC | Acer sp. | 1.34 |
| S10 | ESBC | Betula pendula | 0.27 |
| S10 | ESBC | Carpinus betulus | 3.7 |
| S10 | ESBC | Fagus sylvatica | 53.27 |
| S10 | ESBC | Fraxinus excelsior | 0.4 |
| S10 | ESBC | Larix decidua | 2.78 |
| S10 | ESBC | Picea abies | 0.45 |
| S10 | ESBC | Prunus avium | 0.15 |
| S10 | ESBC | Pseudotsuga menziesii | 0.27 |
| S10 | ESBC | Quercus sp. | 37.3 |
| U01 | Control | Acer sp. | 7.65 |
| U01 | Control | Betula pendula | 0.83 |
| U01 | Control | Carpinus betulus | 5.67 |
| U01 | Control | Fagus sylvatica | 48.17 |
| U01 | Control | Fraxinus excelsior | 14.02 |

| | | | |
|---|---|---|---|
| U01 | Control | Larix decidua | 10.91 |
| U01 | Control | Picea abies | 0.36 |
| U01 | Control | Pinus sylvestris | 1.42 |
| U01 | Control | Prunus avium | 0.03 |
| U01 | Control | Quercus sp. | 10 |
| U01 | Control | Sorbus aucuparia | 0.82 |
| U01 | Control | Tilia sp. | 0.1 |
| U01 | ESBC | Acer sp. | 14.89 |
| U01 | ESBC | Betula pendula | 0.04 |
| U01 | ESBC | Carpinus betulus | 20.73 |
| U01 | ESBC | Fagus sylvatica | 6.94 |
| U01 | ESBC | Fraxinus excelsior | 24.59 |
| U01 | ESBC | Larix decidua | 1.19 |
| U01 | ESBC | Picea abies | 0.18 |
| U01 | ESBC | Pinus sylvestris | 2.79 |
| U01 | ESBC | Prunus avium | 0.14 |
| U01 | ESBC | Quercus sp. | 21.92 |
| U01 | ESBC | Sorbus aucuparia | 5.37 |
| U01 | ESBC | Tilia sp. | 1.16 |
| U01 | ESBC | Ulmus sp. | 0.06 |
| U02 | Control | Acer sp. | 2.95 |
| U02 | Control | Betula pendula | 1.18 |
| U02 | Control | Carpinus betulus | 16.56 |
| U02 | Control | Fagus sylvatica | 22.31 |
| U02 | Control | Fraxinus excelsior | 1.33 |
| U02 | Control | Larix decidua | 18.45 |
| U02 | Control | Picea abies | 3.37 |
| U02 | Control | Pinus sylvestris | 10.12 |
| U02 | Control | Populus tremula | 0.03 |
| U02 | Control | Prunus avium | 1.01 |
| U02 | Control | Quercus sp. | 13.58 |
| U02 | Control | Salix sp. | 0.23 |
| U02 | Control | Sorbus aucuparia | 1.38 |
| U02 | Control | Tilia sp. | 7.51 |
| U02 | ESBC | Acer sp. | 11.84 |
| U02 | ESBC | Aesculus hippocastanum | 0.04 |
| U02 | ESBC | Betula pendula | 0.25 |
| U02 | ESBC | Carpinus betulus | 6.02 |
| U02 | ESBC | Fagus sylvatica | 32.76 |
| U02 | ESBC | Fraxinus excelsior | 19.7 |
| U02 | ESBC | Larix decidua | 5.12 |
| U02 | ESBC | Picea abies | 2.57 |
| U02 | ESBC | Pinus sylvestris | 3.49 |
| U02 | ESBC | Prunus avium | 0.19 |
| U02 | ESBC | Quercus sp. | 12.41 |
| U02 | ESBC | Sorbus aucuparia | 2 |
| U02 | ESBC | Tilia sp. | 3.11 |
| U02 | ESBC | Ulmus sp. | 0.37 |

| | | | |
|---|---|---|---|
| U03 | Control | Acer sp. | 24.56 |
| U03 | Control | Carpinus betulus | 12.98 |
| U03 | Control | Fagus sylvatica | 1.48 |
| U03 | Control | Fraxinus excelsior | 20.27 |
| U03 | Control | Larix decidua | 0.64 |
| U03 | Control | Picea abies | 0.23 |
| U03 | Control | Pinus sylvestris | 14.32 |
| U03 | Control | Prunus avium | 3.38 |
| U03 | Control | Pseudotsuga menziesii | 3.15 |
| U03 | Control | Quercus sp. | 9.42 |
| U03 | Control | Sorbus aucuparia | 3.3 |
| U03 | Control | Tilia sp. | 4.9 |
| U03 | Control | Ulmus sp. | 1.18 |
| U03 | ESBC | Acer sp. | 18.89 |
| U03 | ESBC | Betula pendula | 0.03 |
| U03 | ESBC | Carpinus betulus | 12.99 |
| U03 | ESBC | Fagus sylvatica | 2.28 |
| U03 | ESBC | Fraxinus excelsior | 44.76 |
| U03 | ESBC | Larix decidua | 0.74 |
| U03 | ESBC | Picea abies | 1.05 |
| U03 | ESBC | Pinus sylvestris | 5.72 |
| U03 | ESBC | Populus tremula | 0.14 |
| U03 | ESBC | Prunus avium | 1.35 |
| U03 | ESBC | Pseudotsuga menziesii | 1.97 |
| U03 | ESBC | Quercus sp. | 3.97 |
| U03 | ESBC | Sorbus aucuparia | 2.45 |
| U03 | ESBC | Tilia sp. | 2.73 |
| U03 | ESBC | Ulmus sp. | 0.47 |

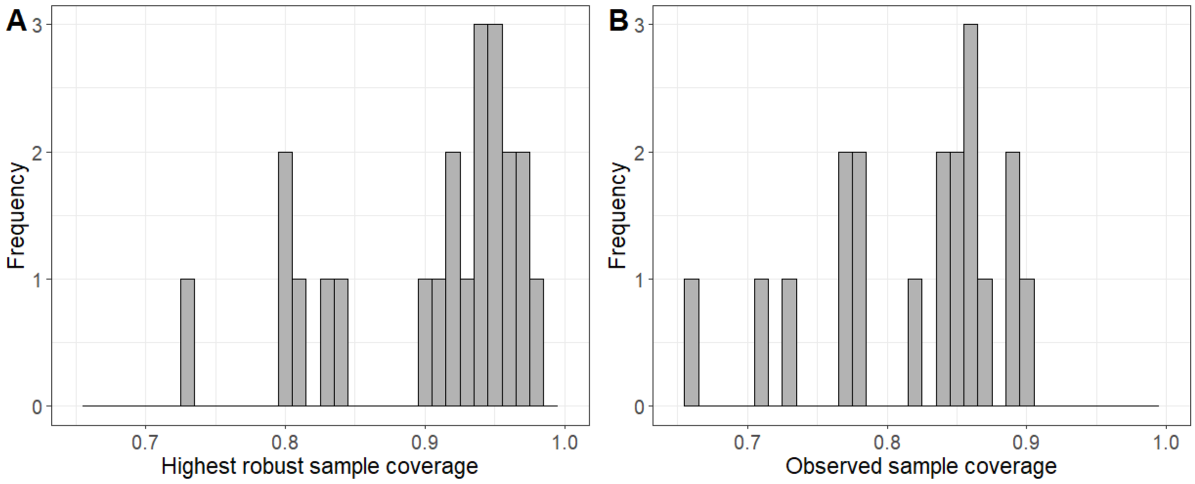


**Figure S1**: Histograms of (A) highest robust sample coverage [0,1] for q = 0 and (B) observed sample coverage for each district (n = 22). The observed distributions illustrate variation in sample completeness across districts.

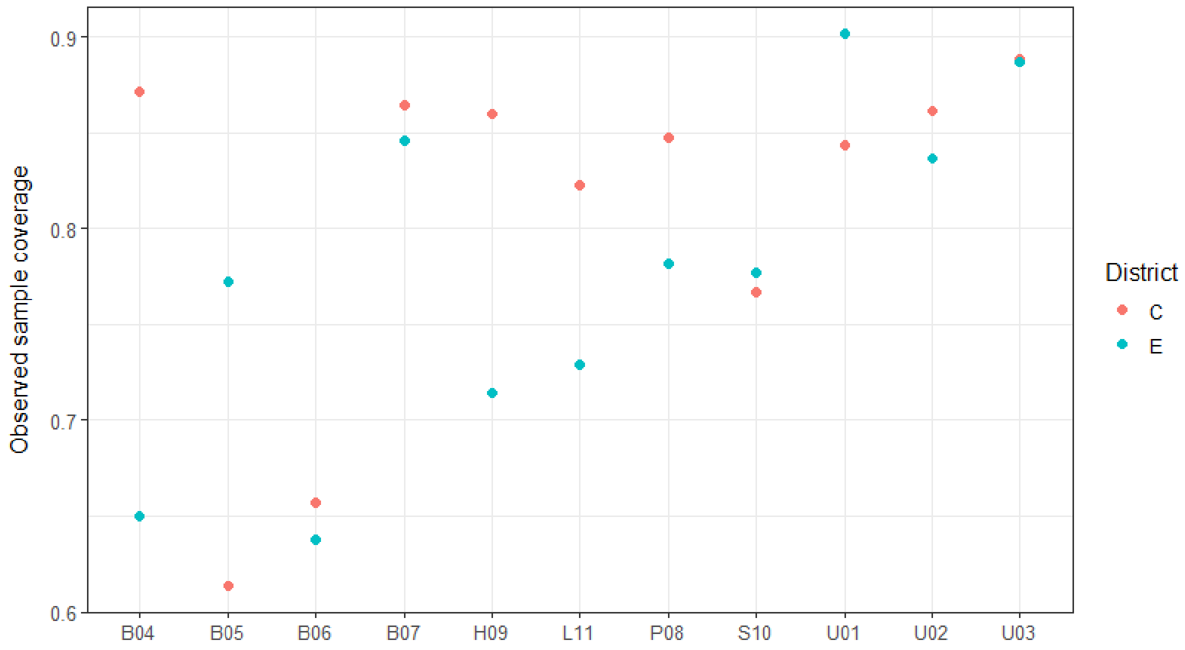


**Figure S2**: Observed sample coverage [0,1] for each district (n = 22) and landscape (n = 11). Values of 1 represent full sample completeness.

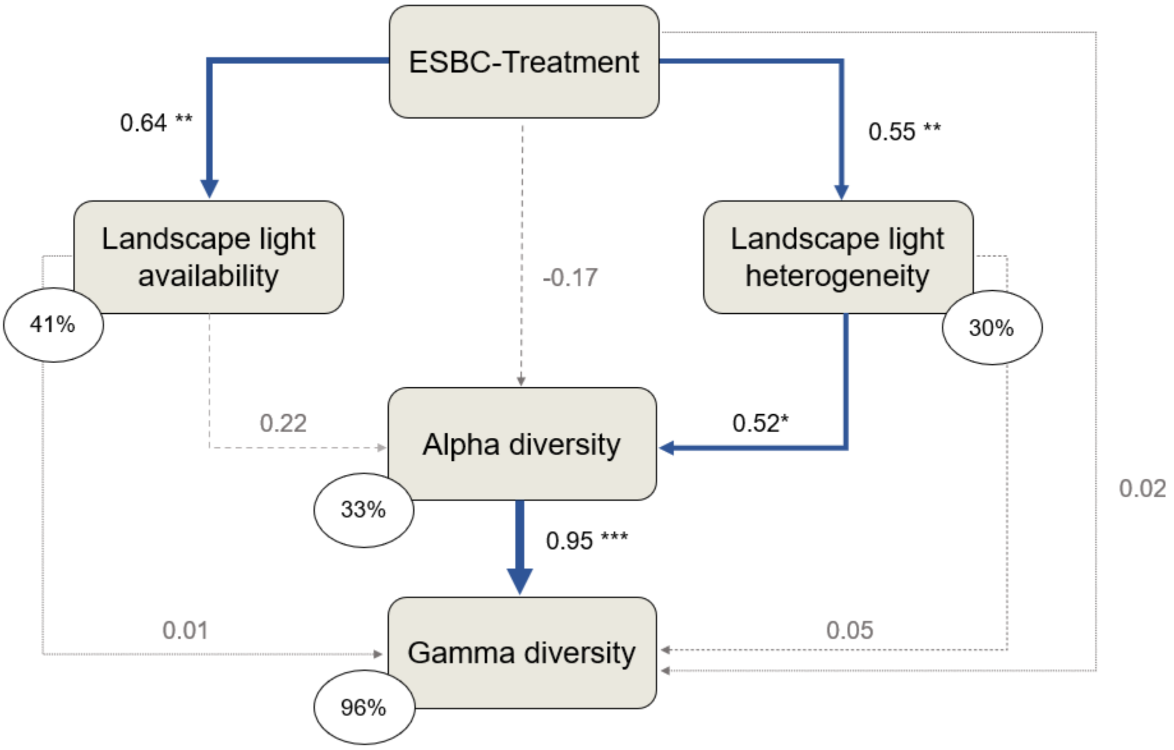


**Figure S3**: **Structural equation model linking understorey plant diversity and management-induced changes in light conditions across multiple spatial scales.** The models were fitted using standardised taxonomic diversity values of the diversity order q=1 for alpha diversity and gamma diversity. Positive relationships between enhanced structural heterogeneity between forest patches (ESBC) and light conditions (light availability and heterogeneity) or plant diversity (mean joint alpha and gamma diversity) indicate a positive ESBC effect. Landscape light availability refers to the mean canopy openness of all patches within a district (mean within-patch canopy openness), while landscape light heterogeneity refers to the coefficient of variation in canopy openness across patches within a district (between-patch canopy openness heterogeneity). Blue arrows indicate significant (* $p < 0.05$, ** $p < 0.01$ and *** $p < 0.001$) and dashed grey arrows non-significant ($p > 0.05$) relationships. Numbers next to arrows are standardised path coefficients and arrow width is proportional to their effect size. Percentage values are explained variances of endogenous variables. The model provided a good fit to the data: Fisher's C = 1.072, df = 2, p-value = 0.585.

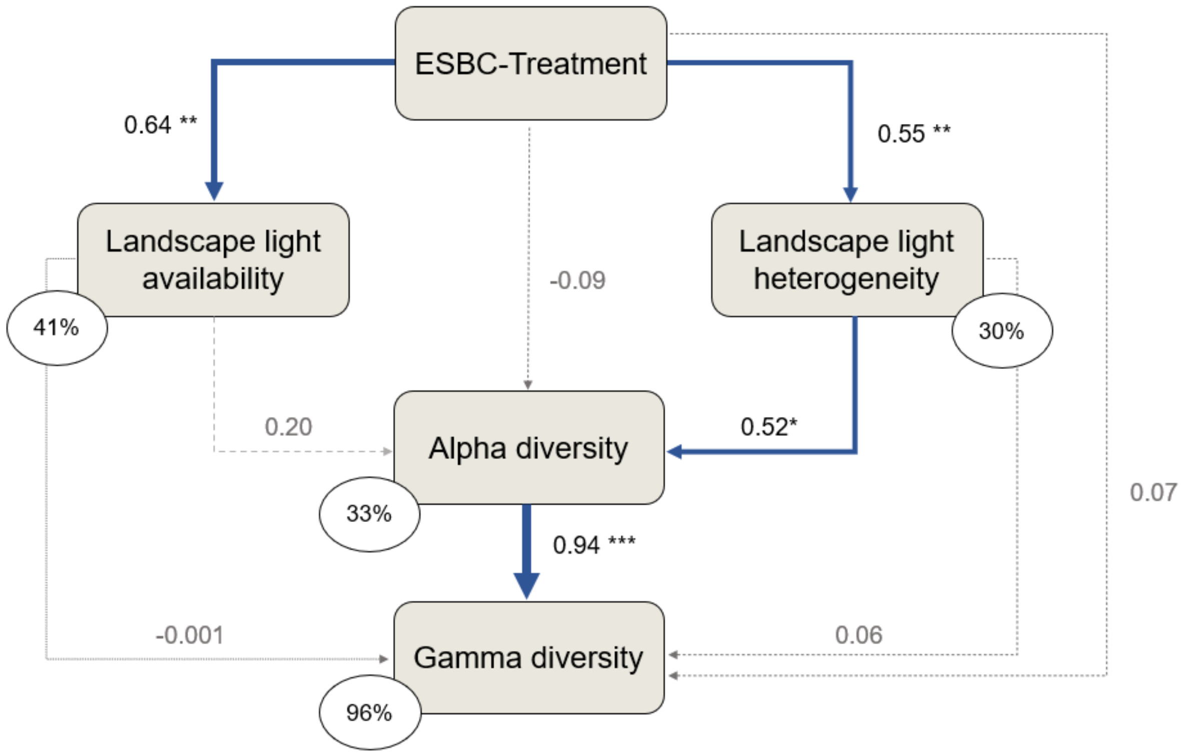


**Figure S4**: **Structural equation model linking understorey plant diversity and management-induced changes in light conditions across multiple spatial scales.** The models were fitted using standardised taxonomic diversity values of the diversity order q=2 for alpha diversity and gamma diversity. For further information see Figure S3. The model provided a good fit to the data: Fisher's C = 1.072, df = 2, p-value = 0.585.

**Gamma with Coverage = 0.9**

| regionSite | TD(q=0) E | TD(q=0) C | Difference | LCL | UCL | W(fixed) |
|---|---|---|---|---|---|---|
| B04 | 23.064 | 17.7 | 5.364 | -5.05 | 15.778 | 12.16% |
| B05 | 36.453 | 16.44 | 20.013 | 0.941 | 39.086 | 3.62% |
| B06 | 57.64 | 10.8 | 46.84 | -4.45 | 98.13 | 0.5% |
| B07 | 27.133 | 6.6 | 20.533 | 10.088 | 30.979 | 12.08% |
| H09 | 38.42 | 8.82 | 29.6 | 1.182 | 58.018 | 1.63% |
| L11 | 59.005 | 22.088 | 36.917 | 14.721 | 59.113 | 2.68% |
| P08 | 41.712 | 18.9 | 22.812 | -1.353 | 46.977 | 2.26% |
| S10 | 45.566 | 17.57 | 27.996 | 18.366 | 37.625 | 14.22% |
| U01 | 82.168 | 69.337 | 12.831 | 2.526 | 23.136 | 12.42% |
| U02 | 82.229 | 57.368 | 24.86 | 17.871 | 31.849 | 26.99% |
| U03 | 103.893 | 86.765 | 17.129 | 6.389 | 27.868 | 11.43% |
| **Meta analysis** | | | **20.323** | **16.692** | **23.955** | |

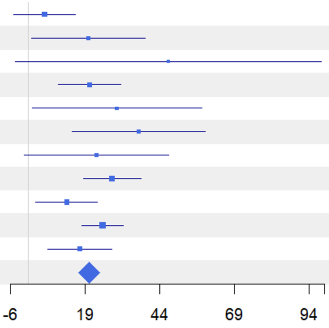


**Figure S5**: Forest plot for taxonomic gamma diversity of order q = 0 showing results of coverage-standardised comparisons between ESBC and C-districts of each individual experimental landscape and across landscapes.

**Alpha with Coverage = 0.9**

| regionSite | TD(q=0) E | TD(q=0) C | Difference | LCL | UCL | W(fixed) |
|---|---|---|---|---|---|---|
| B04 | 18.516 | 5.655 | 12.861 | 0.405 | 25.316 | 0.46% |
| B05 | 12.596 | 7.856 | 4.741 | -0.28 | 9.762 | 2.82% |
| B06 | 15.275 | 4.495 | 10.78 | 4.635 | 16.926 | 1.88% |
| B07 | 8.874 | 2.668 | 6.207 | 4.773 | 7.641 | 34.6% |
| H09 | 11.691 | 3.761 | 7.93 | -0.511 | 16.371 | 1% |
| L11 | 18.512 | 7.152 | 11.359 | 7.458 | 15.26 | 4.67% |
| P08 | 9.617 | 4.489 | 5.128 | 2.82 | 7.436 | 13.35% |
| S10 | 19.521 | 5.783 | 13.739 | 10.928 | 16.55 | 9% |
| U01 | 36.949 | 21.97 | 14.98 | 12.266 | 17.693 | 9.66% |
| U02 | 38.468 | 22.923 | 15.546 | 12.759 | 18.333 | 9.16% |
| U03 | 46.087 | 44.282 | 1.805 | -0.5 | 4.109 | 13.39% |
| **Meta analysis** | | | **8.187** | **7.343** | **9.03** | |

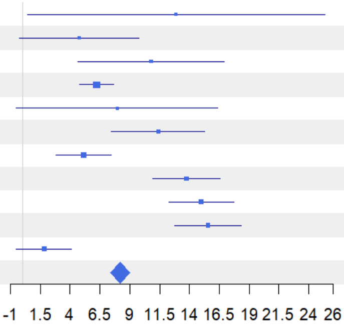


**Figure S6**: Forest plot for taxonomic alpha diversity of order q = 0 showing results of coverage-standardised comparisons between ESBC and C-districts of each individual experimental landscape and across landscapes.

**1-S with Coverage = 0.9**

| regionSite | TD(q=0) E | TD(q=0) C | Difference | LCL | UCL | W(fixed) |
|---|---|---|---|---|---|---|
| B04 | 0.222 | 0.766 | -0.544 | -0.708 | -0.38 | 1.4% |
| B05 | 0.736 | 0.587 | 0.149 | 0.043 | 0.254 | 3.37% |
| B06 | 0.827 | 0.667 | 0.16 | 0.019 | 0.301 | 1.88% |
| B07 | 0.757 | 0.681 | 0.076 | -0.135 | 0.287 | 0.84% |
| H09 | 0.783 | 0.645 | 0.137 | 0.003 | 0.272 | 2.06% |
| L11 | 0.772 | 0.761 | 0.011 | -0.066 | 0.089 | 6.25% |
| P08 | 0.866 | 0.858 | 0.008 | -0.097 | 0.113 | 3.38% |
| S10 | 0.643 | 0.755 | -0.112 | -0.205 | -0.018 | 4.29% |
| U01 | 0.59 | 0.732 | -0.142 | -0.18 | -0.105 | 26.43% |
| U02 | 0.57 | 0.643 | -0.073 | -0.111 | -0.036 | 26.63% |
| U03 | 0.596 | 0.525 | 0.072 | 0.032 | 0.112 | 23.46% |
| **Meta analysis** | | | **-0.04** | **-0.06** | **-0.021** | |

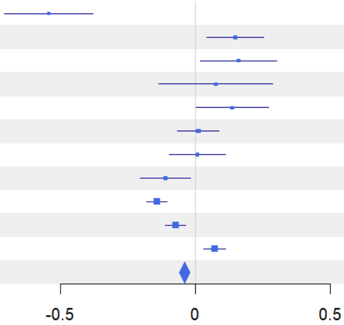


**Figure S7**: Forest plot for taxonomic beta (1-S) diversity of order q = 0 showing results of coverage-standardised comparisons between ESBC and C-districts of each individual experimental landscape and across landscapes.

**Gamma with Coverage = 0.9**

| regionSite | TD(q=1) E | TD(q=1) C | Difference | LCL | UCL | W(fixed) |
|---|---|---|---|---|---|---|
| B04 | 18.543 | 15.041 | 3.502 | -2.639 | 9.643 | 11.76% |
| B05 | 28.431 | 12.75 | 15.681 | 7.863 | 23.499 | 7.25% |
| B06 | 35.735 | 7.36 | 28.375 | 13.587 | 43.162 | 2.03% |
| B07 | 20.298 | 5.705 | 14.593 | 9.051 | 20.134 | 14.44% |
| H09 | 20.96 | 7.489 | 13.471 | 4.146 | 22.796 | 5.1% |
| L11 | 43.747 | 18.601 | 25.146 | 10.873 | 39.419 | 2.18% |
| P08 | 30.718 | 14.322 | 16.396 | 5.509 | 27.282 | 3.74% |
| S10 | 40.761 | 13.91 | 26.852 | 21.87 | 31.833 | 17.86% |
| U01 | 78.245 | 58.921 | 19.324 | 13.301 | 25.347 | 12.22% |
| U02 | 76.958 | 53.725 | 23.233 | 17.885 | 28.581 | 15.5% |
| U03 | 94.685 | 82.274 | 12.411 | 4.938 | 19.884 | 7.94% |
| **Meta analysis** | | | **17.821** | **15.715** | **19.926** | |

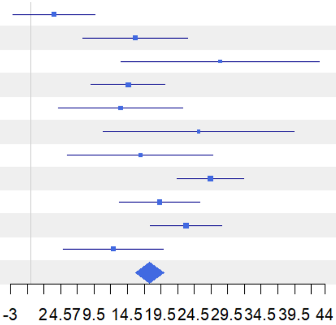


**Figure S8**: Forest plot for taxonomic gamma diversity of order q = 1 showing results of coverage-standardised comparisons between ESBC and C-districts of each individual experimental landscape and across landscapes.

**Alpha with Coverage = 0.9**

| regionSite | TD(q=1) E | TD(q=1) C | Difference | LCL | UCL | W(fixed) |
|---|---|---|---|---|---|---|
| B04 | 9.752 | 4.868 | 4.884 | 1.552 | 8.215 | 2.63% |
| B05 | 9.682 | 6.504 | 3.178 | 0.678 | 5.678 | 4.68% |
| B06 | 13.01 | 3.228 | 9.781 | 7.105 | 12.458 | 4.08% |
| B07 | 7.438 | 2.208 | 5.23 | 4.168 | 6.293 | 25.9% |
| H09 | 6.939 | 2.869 | 4.07 | 1.949 | 6.191 | 6.5% |
| L11 | 12.282 | 5.18 | 7.102 | 4.366 | 9.838 | 3.91% |
| P08 | 7.562 | 3.521 | 4.041 | 2.584 | 5.498 | 13.78% |
| S10 | 15.875 | 4.619 | 11.256 | 9.324 | 13.189 | 7.83% |
| U01 | 31.408 | 16.922 | 14.486 | 12.947 | 16.025 | 12.35% |
| U02 | 30.804 | 18.529 | 12.275 | 10.431 | 14.12 | 8.6% |
| U03 | 38.187 | 36.972 | 1.215 | -0.518 | 2.948 | 9.74% |
| **Meta analysis** | | | **6.975** | **6.434** | **7.515** | |

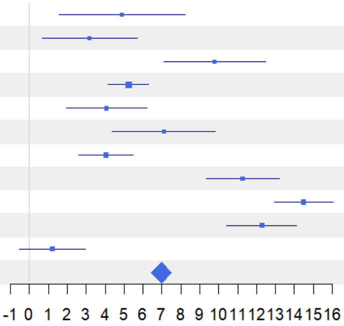


**Figure S9**: Forest plot for taxonomic alpha diversity of order q = 1 showing results of coverage-standardised comparisons between ESBC and C-districts of each individual experimental landscape and across landscapes.

**1-S with Coverage = 0.9**

| regionSite | TD(q=1) E | TD(q=1) C | Difference | LCL | UCL | W(fixed) |
|---|---|---|---|---|---|---|
| B04 | 0.533 | 0.761 | -0.228 | -0.303 | -0.152 | 2.69% |
| B05 | 0.742 | 0.551 | 0.191 | 0.104 | 0.278 | 2.03% |
| B06 | 0.715 | 0.642 | 0.074 | -0.042 | 0.19 | 1.13% |
| B07 | 0.713 | 0.701 | 0.012 | -0.097 | 0.121 | 1.29% |
| H09 | 0.753 | 0.694 | 0.059 | -0.046 | 0.163 | 1.41% |
| L11 | 0.809 | 0.812 | -0.003 | -0.058 | 0.052 | 5.04% |
| P08 | 0.848 | 0.848 | 0 | -0.081 | 0.081 | 2.33% |
| S10 | 0.687 | 0.751 | -0.065 | -0.132 | 0.003 | 3.31% |
| U01 | 0.641 | 0.764 | -0.122 | -0.145 | -0.099 | 28.76% |
| U02 | 0.643 | 0.702 | -0.059 | -0.083 | -0.036 | 27.99% |
| U03 | 0.639 | 0.59 | 0.049 | 0.024 | 0.075 | 24.01% |
| **Meta analysis** | | | **-0.043** | **-0.055** | **-0.03** | |

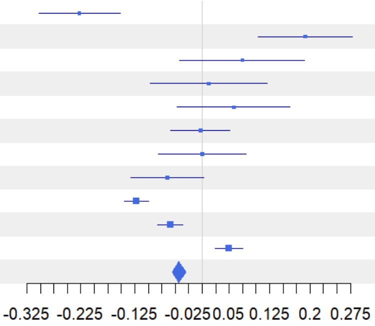


**Figure S10**: Forest plot for taxonomic beta (1-S) diversity of order q = 1 showing results of coverage-standardised comparisons between ESBC and C-districts of each individual experimental landscape and across landscapes.

**Gamma with Coverage = 0.9**

| regionSite | TD(q=2) E | TD(q=2) C | Difference | LCL | UCL | W(fixed) |
|---|---|---|---|---|---|---|
| B04 | 15.476 | 13.548 | 1.928 | -1.853 | 5.709 | 12.33% |
| B05 | 23.414 | 10.595 | 12.819 | 8.374 | 17.264 | 8.92% |
| B06 | 22.331 | 5.651 | 16.68 | 11.694 | 21.666 | 7.09% |
| B07 | 16.726 | 5.161 | 11.564 | 8.204 | 14.925 | 15.6% |
| H09 | 13.54 | 6.78 | 6.76 | 2.169 | 11.351 | 8.36% |
| L11 | 34.337 | 16.17 | 18.167 | 13.203 | 23.131 | 7.15% |
| P08 | 23.439 | 11.163 | 12.276 | 6.482 | 18.069 | 5.25% |
| S10 | 37.377 | 11.47 | 25.907 | 22.042 | 29.772 | 11.8% |
| U01 | 75.461 | 52.826 | 22.636 | 17.947 | 27.324 | 8.02% |
| U02 | 73.323 | 51.208 | 22.115 | 17.922 | 26.309 | 10.02% |
| U03 | 89.053 | 79.269 | 9.784 | 4.107 | 15.461 | 5.47% |
| **Meta analysis** | | | **14.498** | **13.171** | **15.826** | |

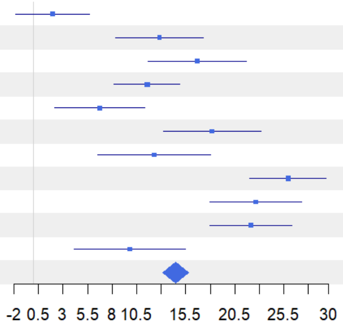


**Figure S11**: Forest plot for taxonomic gamma diversity of order q = 2 showing results of coverage-standardised comparisons between ESBC and C-districts of each individual experimental landscape and across landscapes.

**Alpha with Coverage = 0.9**

| regionSite | TD(q=2) E | TD(q=2) C | Difference | LCL | UCL | W(fixed) |
|---|---|---|---|---|---|---|
| B04 | 5.001 | 4.233 | 0.768 | -0.371 | 1.907 | 7.93% |
| B05 | 7.442 | 4.616 | 2.826 | 1.535 | 4.118 | 6.16% |
| B06 | 10.095 | 2.091 | 8.004 | 6.233 | 9.775 | 3.28% |
| B07 | 6.369 | 1.874 | 4.496 | 3.695 | 5.296 | 16.06% |
| H09 | 4.244 | 2.388 | 1.856 | 0.931 | 2.781 | 12.02% |
| L11 | 8.187 | 4.042 | 4.145 | 3.08 | 5.209 | 9.07% |
| P08 | 5.842 | 2.899 | 2.943 | 2.166 | 3.719 | 17.04% |
| S10 | 12.51 | 3.568 | 8.943 | 7.676 | 10.209 | 6.41% |
| U01 | 27.927 | 13.867 | 14.06 | 13.043 | 15.077 | 9.94% |
| U02 | 25.621 | 15.698 | 9.923 | 8.748 | 11.098 | 7.45% |
| U03 | 33.481 | 32.536 | 0.945 | -0.542 | 2.433 | 4.65% |
| **Meta analysis** | | | **5.074** | **4.753** | **5.394** | |

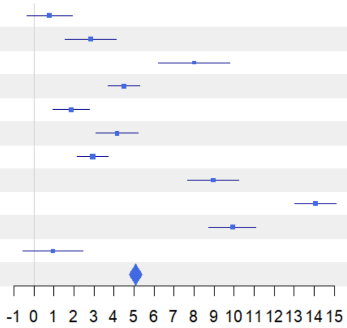


**Figure S12**: Forest plot for taxonomic alpha diversity of order q = 2 showing results of coverage-standardised comparisons between ESBC and C-districts of each individual experimental landscape and across landscapes.

**1-S with Coverage = 0.9**

| regionSite | TD(q=2) E | TD(q=2) C | Difference | LCL | UCL | W(fixed) |
|---|---|---|---|---|---|---|
| B04 | 0.761 | 0.774 | -0.012 | -0.058 | 0.034 | 4.07% |
| B05 | 0.767 | 0.635 | 0.133 | 0.085 | 0.18 | 3.85% |
| B06 | 0.616 | 0.72 | -0.104 | -0.168 | -0.039 | 2.05% |
| B07 | 0.697 | 0.728 | -0.031 | -0.133 | 0.07 | 0.84% |
| H09 | 0.772 | 0.729 | 0.044 | -0.033 | 0.12 | 1.46% |
| L11 | 0.857 | 0.844 | 0.013 | -0.025 | 0.051 | 5.84% |
| P08 | 0.845 | 0.833 | 0.012 | -0.054 | 0.077 | 2.01% |
| S10 | 0.748 | 0.775 | -0.027 | -0.075 | 0.022 | 3.68% |
| U01 | 0.675 | 0.79 | -0.115 | -0.134 | -0.097 | 24.79% |
| U02 | 0.697 | 0.743 | -0.046 | -0.063 | -0.029 | 28.66% |
| U03 | 0.669 | 0.632 | 0.037 | 0.017 | 0.056 | 22.74% |
| **Meta analysis** | | | **-0.03** | **-0.04** | **-0.021** | |

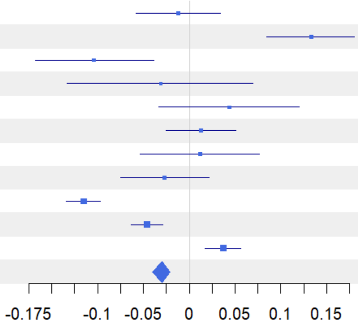


**Figure S13**: Forest plot for taxonomic beta (1-S) diversity of order q = 2 showing results of coverage-standardised comparisons between ESBC and C-districts of each individual experimental landscape and across landscapes.

**Gamma with Coverage = 0.9**

| regionSite | PD(q=0) E | PD(q=0) C | Difference | LCL | UCL | W(fixed) |
|---|---|---|---|---|---|---|
| B04 | 6.82 | 6.299 | 0.521 | -4.829 | 5.871 | 7.07% |
| B05 | 8.953 | 5.569 | 3.383 | -3.509 | 10.275 | 4.26% |
| B06 | 10.981 | 4.203 | 6.778 | 1.453 | 12.103 | 7.14% |
| B07 | 7.717 | 3.53 | 4.187 | 0.94 | 7.434 | 19.2% |
| H09 | 7.757 | 4.288 | 3.469 | -5.366 | 12.303 | 2.59% |
| L11 | 11.101 | 6.21 | 4.891 | -1.007 | 10.79 | 5.82% |
| P08 | 7.987 | 5.836 | 2.151 | -2.75 | 7.052 | 8.43% |
| S10 | 9.252 | 5.892 | 3.361 | -2.735 | 9.457 | 5.45% |
| U01 | 13.62 | 13.281 | 0.339 | -3.938 | 4.616 | 11.06% |
| U02 | 13.503 | 11.229 | 2.275 | -2.259 | 6.808 | 9.85% |
| U03 | 16.353 | 15.054 | 1.299 | -1.954 | 4.553 | 19.13% |
| **Meta analysis** | | | **2.718** | **1.295** | **4.141** | |

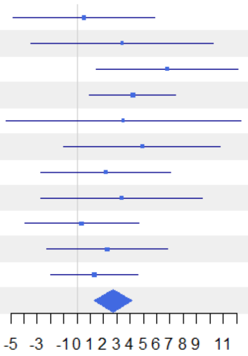


**Figure S14**: Forest plot for phylogenetic gamma diversity of order q = 0 showing results of coverage-standardised comparisons between ESBC and C-districts of each individual experimental landscape and across landscapes.

**Alpha with Coverage = 0.9**

| regionSite | PD(q=0) E | PD(q=0) C | Difference | LCL | UCL | W(fixed) |
|---|---|---|---|---|---|---|
| B04 | 5.509 | 2.914 | 2.594 | 1.527 | 3.661 | 6.36% |
| B05 | 4.757 | 3.744 | 1.013 | -0.338 | 2.363 | 3.97% |
| B06 | 5.237 | 1.945 | 3.292 | 2.114 | 4.469 | 5.22% |
| B07 | 4.245 | 1.794 | 2.451 | 1.748 | 3.154 | 14.66% |
| H09 | 4.212 | 2.089 | 2.123 | 0.99 | 3.255 | 5.65% |
| L11 | 4.093 | 2.58 | 1.513 | 0.514 | 2.511 | 7.27% |
| P08 | 3.824 | 2.324 | 1.5 | 0.67 | 2.33 | 10.51% |
| S10 | 5.473 | 2.464 | 3.009 | 2.007 | 4.011 | 7.22% |
| U01 | 7.404 | 5.535 | 1.868 | 1.192 | 2.545 | 15.81% |
| U02 | 8.052 | 5.787 | 2.265 | 1.519 | 3.012 | 13% |
| U03 | 9.462 | 9.603 | -0.14 | -0.977 | 0.696 | 10.35% |
| **Meta analysis** | | | **1.916** | **1.647** | **2.185** | |

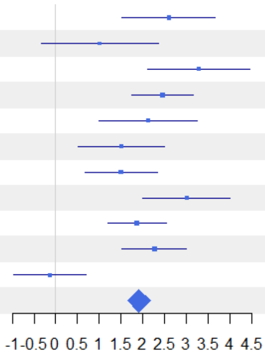


**Figure S15**: Forest plot for phylogenetic alpha diversity of order q = 0 showing results of coverage-standardised comparisons between ESBC and C-districts of each individual experimental landscape and across landscapes.

**1-S with Coverage = 0.9**

| regionSite | PD(q=0) E | PD(q=0) C | Difference | LCL | UCL | W(fixed) |
|---|---|---|---|---|---|---|
| B04 | 0.216 | 0.604 | -0.388 | -0.571 | -0.205 | 3.66% |
| B05 | 0.527 | 0.369 | 0.159 | -0.059 | 0.376 | 2.59% |
| B06 | 0.589 | 0.604 | -0.016 | -0.205 | 0.174 | 3.41% |
| B07 | 0.506 | 0.553 | -0.047 | -0.218 | 0.124 | 4.19% |
| H09 | 0.514 | 0.577 | -0.063 | -0.275 | 0.149 | 2.72% |
| L11 | 0.71 | 0.658 | 0.053 | -0.065 | 0.17 | 8.91% |
| P08 | 0.586 | 0.677 | -0.091 | -0.24 | 0.058 | 5.52% |
| S10 | 0.459 | 0.654 | -0.195 | -0.322 | -0.068 | 7.56% |
| U01 | 0.489 | 0.625 | -0.136 | -0.25 | -0.022 | 9.43% |
| U02 | 0.433 | 0.519 | -0.087 | -0.18 | 0.006 | 14.2% |
| U03 | 0.451 | 0.388 | 0.063 | 0.007 | 0.12 | 37.8% |
| **Meta analysis** | | | **-0.031** | **-0.065** | **0.004** | |

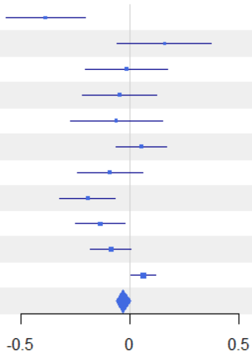


**Figure S16**: Forest plot for phylogenetic beta (1-S) diversity of order q = 0 showing results of coverage-standardised comparisons between ESBC and C-districts of each individual experimental landscape and across landscapes.

**Gamma with Coverage = 0.9**

| regionSite | PD(q=1) E | PD(q=1) C | Difference | LCL | UCL | W(fixed) |
|---|---|---|---|---|---|---|
| B04 | 5.616 | 5.626 | -0.009 | -2.345 | 2.326 | 10.84% |
| B05 | 7.735 | 4.938 | 2.797 | -0.054 | 5.649 | 7.27% |
| B06 | 6.89 | 3.173 | 3.717 | 1.356 | 6.079 | 10.6% |
| B07 | 6.158 | 3.282 | 2.876 | 1.353 | 4.399 | 25.49% |
| H09 | 5.266 | 3.627 | 1.639 | -3.407 | 6.685 | 2.32% |
| L11 | 8.524 | 4.98 | 3.545 | 0.481 | 6.609 | 6.3% |
| P08 | 6.95 | 4.918 | 2.033 | -0.31 | 4.375 | 10.77% |
| S10 | 8.543 | 4.623 | 3.92 | 0.212 | 7.627 | 4.3% |
| U01 | 13.072 | 11.917 | 1.155 | -2.082 | 4.391 | 5.64% |
| U02 | 12.902 | 10.678 | 2.224 | -1.072 | 5.52 | 5.44% |
| U03 | 15.391 | 14.442 | 0.949 | -1.365 | 3.263 | 11.04% |
| **Meta analysis** | | | **2.269** | **1.5** | **3.038** | |

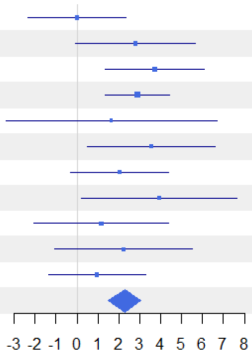


**Figure S17**: Forest plot for phylogenetic gamma diversity of order q = 1 showing results of coverage-standardised comparisons between ESBC and C-districts of each individual experimental landscape and across landscapes.

**Alpha with Coverage = 0.9**

| regionSite | PD(q=1) E | PD(q=1) C | Difference | LCL | UCL | W(fixed) |
|---|---|---|---|---|---|---|
| B04 | 3.66 | 2.537 | 1.123 | 0.659 | 1.586 | 8.52% |
| B05 | 4.016 | 2.902 | 1.114 | 0.469 | 1.759 | 4.4% |
| B06 | 4.356 | 1.562 | 2.794 | 2.268 | 3.32 | 6.62% |
| B07 | 3.752 | 1.472 | 2.28 | 1.89 | 2.671 | 12% |
| H09 | 2.517 | 1.706 | 0.811 | 0.371 | 1.252 | 9.43% |
| L11 | 2.96 | 2.022 | 0.938 | 0.514 | 1.361 | 10.22% |
| P08 | 3.231 | 1.975 | 1.256 | 0.876 | 1.636 | 12.69% |
| S10 | 4.387 | 1.888 | 2.499 | 2.021 | 2.977 | 8.03% |
| U01 | 6.485 | 4.459 | 2.026 | 1.632 | 2.421 | 11.79% |
| U02 | 6.584 | 4.8 | 1.784 | 1.352 | 2.215 | 9.84% |
| U03 | 8.066 | 8.157 | -0.091 | -0.624 | 0.441 | 6.46% |
| **Meta analysis** | | | **1.544** | **1.409** | **1.68** | |

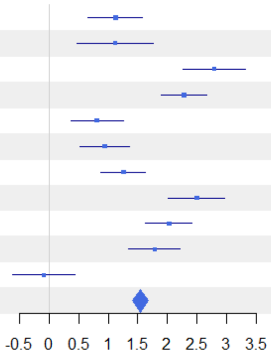


**Figure S18**: Forest plot for phylogenetic alpha diversity of order q = 1 showing results of coverage-standardised comparisons between ESBC and C-districts of each individual experimental landscape and across landscapes.

**1-S with Coverage = 0.9**

| regionSite | PD(q=1) E | PD(q=1) C | Difference | LCL | UCL | W(fixed) |
|---|---|---|---|---|---|---|
| B04 | 0.392 | 0.618 | -0.226 | -0.341 | -0.11 | 5.19% |
| B05 | 0.541 | 0.464 | 0.077 | -0.088 | 0.242 | 2.54% |
| B06 | 0.414 | 0.571 | -0.157 | -0.276 | -0.039 | 4.91% |
| B07 | 0.44 | 0.621 | -0.181 | -0.287 | -0.075 | 6.17% |
| H09 | 0.587 | 0.596 | -0.009 | -0.164 | 0.147 | 2.86% |
| L11 | 0.734 | 0.668 | 0.066 | -0.029 | 0.161 | 7.62% |
| P08 | 0.602 | 0.673 | -0.071 | -0.174 | 0.032 | 6.53% |
| S10 | 0.547 | 0.666 | -0.118 | -0.223 | -0.014 | 6.3% |
| U01 | 0.54 | 0.671 | -0.131 | -0.232 | -0.029 | 6.68% |
| U02 | 0.525 | 0.59 | -0.065 | -0.148 | 0.018 | 10.07% |
| U03 | 0.51 | 0.466 | 0.044 | 0.003 | 0.085 | 41.13% |
| **Meta analysis** | | | **-0.033** | **-0.06** | **-0.007** | |

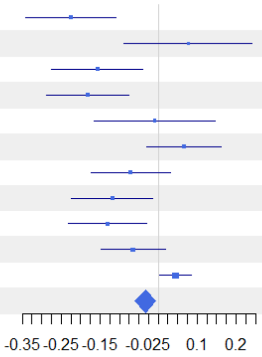


**Figure S19**: Forest plot for phylogenetic beta (1-S) diversity of order q = 1 showing results of coverage-standardised comparisons between ESBC and C-districts of each individual experimental landscape and across landscapes.

**Gamma with Coverage = 0.9**

| regionSite | PD(q=2) E | PD(q=2) C | Difference | LCL | UCL | W(fixed) |
|---|---|---|---|---|---|---|
| B04 | 5.01 | 5.272 | -0.262 | -1.861 | 1.337 | 11.18% |
| B05 | 7.128 | 4.566 | 2.562 | 0.377 | 4.747 | 5.99% |
| B06 | 5.636 | 2.772 | 2.864 | 1.48 | 4.248 | 14.92% |
| B07 | 5.518 | 3.149 | 2.37 | 1.289 | 3.45 | 24.49% |
| H09 | 4.149 | 3.326 | 0.823 | -1.698 | 3.344 | 4.5% |
| L11 | 7.368 | 4.31 | 3.058 | 1.153 | 4.963 | 7.88% |
| P08 | 6.319 | 4.4 | 1.919 | -0.078 | 3.916 | 7.17% |
| S10 | 8.121 | 3.875 | 4.245 | 2.504 | 5.986 | 9.43% |
| U01 | 12.7 | 11.188 | 1.512 | -1.411 | 4.436 | 3.34% |
| U02 | 12.505 | 10.32 | 2.186 | -0.679 | 5.05 | 3.48% |
| U03 | 14.826 | 14.046 | 0.78 | -1.158 | 2.717 | 7.61% |
| **Meta analysis** | | | **2.134** | **1.599** | **2.668** | |

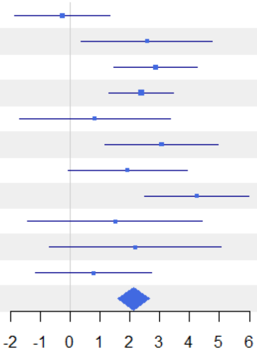


**Figure S20**: Forest plot for phylogenetic gamma diversity of order q = 2 showing results of coverage-standardised comparisons between ESBC and C-districts of each individual experimental landscape and across landscapes.

**Alpha with Coverage = 0.9**

| regionSite | PD(q=2) E | PD(q=2) C | Difference | LCL | UCL | W(fixed) |
|---|---|---|---|---|---|---|
| B04 | 2.792 | 2.291 | 0.501 | 0.191 | 0.811 | 9.01% |
| B05 | 3.383 | 2.27 | 1.113 | 0.712 | 1.514 | 5.37% |
| B06 | 3.578 | 1.171 | 2.407 | 2.081 | 2.733 | 8.16% |
| B07 | 3.438 | 1.265 | 2.173 | 1.865 | 2.481 | 9.12% |
| H09 | 1.953 | 1.535 | 0.418 | 0.16 | 0.675 | 13.07% |
| L11 | 2.481 | 1.78 | 0.701 | 0.446 | 0.956 | 13.31% |
| P08 | 2.792 | 1.759 | 1.033 | 0.734 | 1.331 | 9.74% |
| S10 | 3.673 | 1.586 | 2.087 | 1.765 | 2.41 | 8.34% |
| U01 | 5.974 | 3.922 | 2.052 | 1.761 | 2.342 | 10.26% |
| U02 | 5.761 | 4.282 | 1.479 | 1.159 | 1.8 | 8.42% |
| U03 | 7.319 | 7.385 | -0.066 | -0.474 | 0.342 | 5.2% |
| **Meta analysis** | | | **1.254** | **1.161** | **1.347** | |

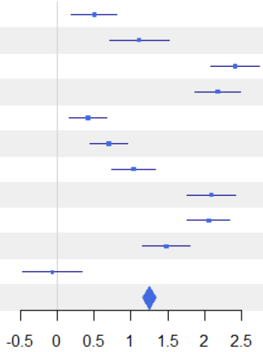


**Figure S21**: Forest plot for phylogenetic alpha diversity of order q = 2 showing results of coverage-standardised comparisons between ESBC and C-districts of each individual experimental landscape and across landscapes.

**1-S with Coverage = 0.9**

| regionSite | PD(q=2) E | PD(q=2) C | Difference | LCL | UCL | W(fixed) |
|---|---|---|---|---|---|---|
| B04 | 0.498 | 0.636 | -0.138 | -0.254 | -0.023 | 3.99% |
| B05 | 0.591 | 0.566 | 0.025 | -0.156 | 0.207 | 1.61% |
| B06 | 0.411 | 0.65 | -0.239 | -0.324 | -0.154 | 7.32% |
| B07 | 0.424 | 0.673 | -0.249 | -0.338 | -0.16 | 6.74% |
| H09 | 0.596 | 0.606 | -0.01 | -0.159 | 0.139 | 2.4% |
| L11 | 0.746 | 0.66 | 0.086 | -0.006 | 0.178 | 6.3% |
| P08 | 0.628 | 0.675 | -0.047 | -0.166 | 0.072 | 3.75% |
| S10 | 0.616 | 0.665 | -0.048 | -0.129 | 0.032 | 8.22% |
| U01 | 0.567 | 0.696 | -0.128 | -0.228 | -0.029 | 5.33% |
| U02 | 0.578 | 0.627 | -0.049 | -0.133 | 0.034 | 7.62% |
| U03 | 0.543 | 0.508 | 0.034 | 0.001 | 0.068 | 46.72% |
| **Meta analysis** | | | **-0.034** | **-0.058** | **-0.011** | |

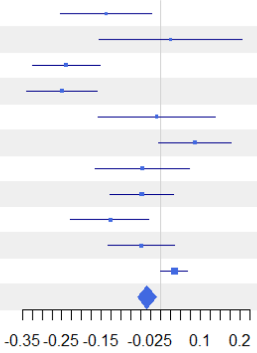


**Figure S22**: Forest plot for phylogenetic beta (1-S) diversity of order q = 2 showing results of coverage-standardised comparisons between ESBC and C-districts of each individual experimental landscape and across landscapes.

**Gamma with Coverage = 0.9**

| regionSite | FD(q=0) E | FD(q=0) C | Difference | LCL | UCL | W(fixed) |
|---|---|---|---|---|---|---|
| B04 | 2.448 | 2.134 | 0.314 | -1.112 | 1.74 | 3.97% |
| B05 | 2.834 | 2.043 | 0.791 | -1.1 | 2.682 | 2.25% |
| B06 | 2.596 | 1.944 | 0.652 | -1.441 | 2.746 | 1.84% |
| B07 | 2.27 | 1.929 | 0.341 | -0.477 | 1.159 | 12.04% |
| H09 | 2.379 | 1.625 | 0.754 | -1.114 | 2.622 | 2.31% |
| L11 | 3.173 | 2.223 | 0.95 | -0.777 | 2.677 | 2.7% |
| P08 | 3.193 | 2.115 | 1.079 | -0.281 | 2.438 | 4.36% |
| S10 | 3.431 | 1.929 | 1.503 | 0.469 | 2.536 | 7.54% |
| U01 | 3.401 | 2.849 | 0.552 | -0.124 | 1.228 | 17.65% |
| U02 | 3.317 | 2.855 | 0.462 | -0.214 | 1.138 | 17.62% |
| U03 | 3.651 | 3.47 | 0.18 | -0.359 | 0.719 | 27.71% |
| **Meta analysis** | | | **0.516** | **0.232** | **0.799** | |

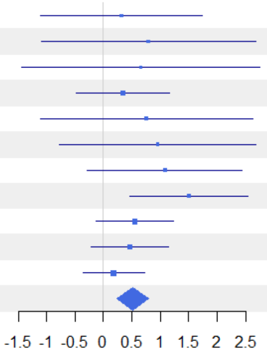


**Figure S23**: Forest plot for functional gamma diversity of order q = 0 showing results of coverage-standardised comparisons between ESBC and C-districts of each individual experimental landscape and across landscapes.

**Alpha with Coverage = 0.9**

| regionSite | FD(q=0) E | FD(q=0) C | Difference | LCL | UCL | W(fixed) |
|---|---|---|---|---|---|---|
| B04 | 2.836 | 1.816 | 1.019 | 0.373 | 1.666 | 3.54% |
| B05 | 2.521 | 2.303 | 0.217 | -0.437 | 0.871 | 3.47% |
| B06 | 2.621 | 2.109 | 0.512 | -0.26 | 1.284 | 2.49% |
| B07 | 2.159 | 1.789 | 0.37 | -0.037 | 0.776 | 8.98% |
| H09 | 2.416 | 1.507 | 0.909 | 0.36 | 1.458 | 4.92% |
| L11 | 2.65 | 1.954 | 0.696 | 0.192 | 1.2 | 5.83% |
| P08 | 2.365 | 1.7 | 0.665 | 0.159 | 1.17 | 5.8% |
| S10 | 2.819 | 1.711 | 1.108 | 0.637 | 1.58 | 6.66% |
| U01 | 3.002 | 2.429 | 0.573 | 0.316 | 0.83 | 22.46% |
| U02 | 2.966 | 2.409 | 0.557 | 0.24 | 0.874 | 14.77% |
| U03 | 3.338 | 3.257 | 0.081 | -0.184 | 0.346 | 21.08% |
| **Meta analysis** | | | **0.515** | **0.393** | **0.637** | |

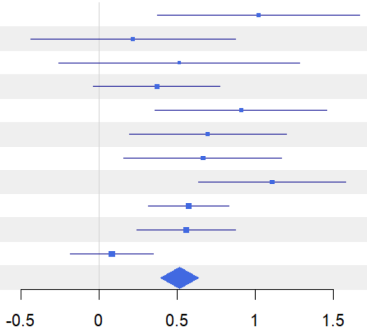


**Figure S24**: Forest plot for functional alpha diversity of order q = 0 showing results of coverage-standardised comparisons between ESBC and C-districts of each individual experimental landscape and across landscapes.

**1-S with Coverage = 0.9**

| regionSite | FD(q=0) E | FD(q=0) C | Difference | LCL | UCL | W(fixed) |
|---|---|---|---|---|---|---|
| B04 | -0.041 | 0.065 | -0.106 | -0.345 | 0.132 | 2.01% |
| B05 | 0.022 | -0.197 | 0.219 | -0.003 | 0.441 | 2.32% |
| B06 | -0.01 | -0.153 | 0.143 | -0.102 | 0.388 | 1.9% |
| B07 | 0.014 | 0.037 | -0.023 | -0.203 | 0.156 | 3.53% |
| H09 | -0.019 | 0.05 | -0.068 | -0.27 | 0.134 | 2.79% |
| L11 | 0.01 | 0.046 | -0.035 | -0.167 | 0.097 | 6.56% |
| P08 | 0.109 | 0.104 | 0.005 | -0.163 | 0.173 | 4.05% |
| S10 | 0.007 | 0.033 | -0.027 | -0.209 | 0.156 | 3.42% |
| U01 | 0.021 | 0.019 | 0.002 | -0.067 | 0.071 | 24.03% |
| U02 | 0.013 | 0.026 | -0.013 | -0.093 | 0.067 | 17.76% |
| U03 | 0.004 | -0.007 | 0.01 | -0.05 | 0.07 | 31.62% |
| **Meta analysis** | | | **0.001** | **-0.032** | **0.035** | |

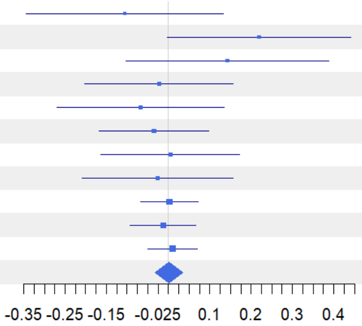


**Figure S25**: Forest plot for functional beta (1-S) diversity of order q = 0 showing results of coverage-standardised comparisons between ESBC and C-districts of each individual experimental landscape and across landscapes.

Gamma with Coverage = 0.9

| regionSite | FD(q=1) E | FD(q=1) C | Difference | LCL | UCL | W(fixed) |
|---|---|---|---|---|---|---|
| B04 | 1.981 | 1.781 | 0.2 | -0.11 | 0.51 | 6.71% |
| B05 | 2.192 | 1.665 | 0.527 | 0.075 | 0.98 | 3.15% |
| B06 | 2.211 | 1.772 | 0.439 | -0.006 | 0.884 | 3.26% |
| B07 | 1.927 | 1.789 | 0.138 | -0.121 | 0.397 | 9.62% |
| H09 | 1.691 | 1.367 | 0.324 | -0.054 | 0.702 | 4.51% |
| L11 | 2.101 | 1.619 | 0.482 | 0.117 | 0.847 | 4.84% |
| P08 | 2.221 | 1.645 | 0.576 | 0.236 | 0.917 | 5.57% |
| S10 | 2.433 | 1.552 | 0.881 | 0.534 | 1.228 | 5.36% |
| U01 | 2.664 | 2.274 | 0.39 | 0.209 | 0.571 | 19.72% |
| U02 | 2.593 | 2.326 | 0.267 | 0.075 | 0.459 | 17.54% |
| U03 | 2.794 | 2.707 | 0.087 | -0.094 | 0.268 | 19.71% |
| **Meta analysis** | | | **0.316** | **0.235** | **0.396** | |

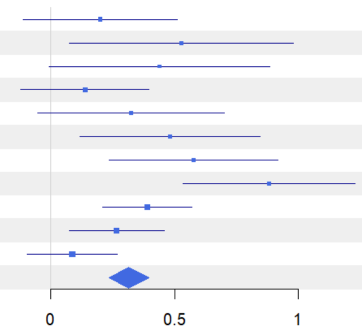


**Figure S26**: Forest plot for functional gamma diversity of order q = 1 showing results of coverage-standardised comparisons between ESBC and C-districts of each individual experimental landscape and across landscapes.

Alpha with Coverage = 0.9

| regionSite | FD(q=1) E | FD(q=1) C | Difference | LCL | UCL | W(fixed) |
|---|---|---|---|---|---|---|
| B04 | 2.147 | 1.511 | 0.635 | 0.327 | 0.944 | 3.07% |
| B05 | 2.006 | 1.675 | 0.331 | -0.12 | 0.781 | 1.44% |
| B06 | 2.197 | 1.484 | 0.713 | 0.261 | 1.165 | 1.43% |
| B07 | 1.924 | 1.461 | 0.462 | 0.216 | 0.709 | 4.82% |
| H09 | 1.75 | 1.333 | 0.416 | 0.14 | 0.692 | 3.84% |
| L11 | 1.549 | 1.558 | -0.009 | -0.248 | 0.23 | 5.13% |
| P08 | 1.96 | 1.5 | 0.46 | 0.219 | 0.701 | 5.03% |
| S10 | 1.893 | 1.468 | 0.425 | 0.189 | 0.661 | 5.27% |
| U01 | 2.466 | 1.959 | 0.507 | 0.402 | 0.612 | 26.68% |
| U02 | 2.435 | 1.9 | 0.535 | 0.418 | 0.653 | 21.15% |
| U03 | 2.649 | 2.611 | 0.038 | -0.077 | 0.153 | 22.14% |
| **Meta analysis** | | | **0.375** | **0.321** | **0.429** | |

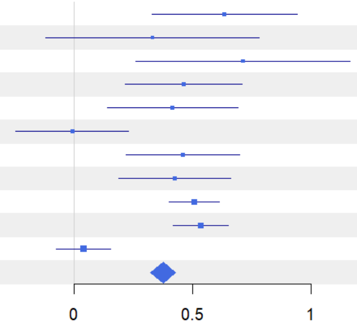


**Figure S27**: Forest plot for functional alpha diversity of order q = 1 showing results of coverage-standardised comparisons between ESBC and C-districts of each individual experimental landscape and across landscapes.

1-S with Coverage = 0.9

| regionSite | FD(q=1) E | FD(q=1) C | Difference | LCL | UCL | W(fixed) |
|---|---|---|---|---|---|---|
| B04 | 0.001 | 0.113 | -0.113 | -0.229 | 0.003 | 2.34% |
| B05 | 0.091 | 0.038 | 0.053 | -0.082 | 0.189 | 1.71% |
| B06 | 0.069 | 0.186 | -0.117 | -0.273 | 0.039 | 1.29% |
| B07 | 0.006 | 0.194 | -0.188 | -0.285 | -0.091 | 3.35% |
| H09 | -0.005 | 0.026 | -0.031 | -0.137 | 0.075 | 2.79% |
| L11 | 0.283 | 0.045 | 0.238 | 0.167 | 0.309 | 6.26% |
| P08 | 0.102 | 0.069 | 0.033 | -0.074 | 0.141 | 2.71% |
| S10 | 0.21 | 0.064 | 0.146 | 0.051 | 0.241 | 3.46% |
| U01 | 0.025 | 0.099 | -0.074 | -0.113 | -0.035 | 20.87% |
| U02 | 0.017 | 0.144 | -0.127 | -0.166 | -0.088 | 21.01% |
| U03 | 0.014 | 0.011 | 0.003 | -0.027 | 0.033 | 34.23% |
| **Meta analysis** | | | **-0.031** | **-0.048** | **-0.013** | |

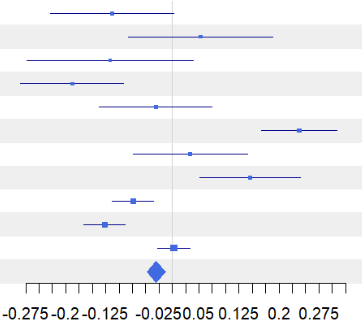


**Figure S28**: Forest plot for functional beta (1-S) diversity of order q = 1 showing results of coverage-standardised comparisons between ESBC and C-districts of each individual experimental landscape and across landscapes.

**Gamma with Coverage = 0.9**

| regionSite | FD(q=2) E | FD(q=2) C | Difference | LCL | UCL | W(fixed) |
|---|---|---|---|---|---|---|
| B04 | 1.821 | 1.619 | 0.202 | 0.012 | 0.391 | 8.23% |
| B05 | 1.932 | 1.518 | 0.415 | 0.124 | 0.705 | 3.49% |
| B06 | 2.028 | 1.69 | 0.338 | 0.068 | 0.608 | 4.05% |
| B07 | 1.782 | 1.709 | 0.073 | -0.131 | 0.276 | 7.09% |
| H09 | 1.493 | 1.276 | 0.217 | -0.004 | 0.437 | 6.06% |
| L11 | 1.756 | 1.446 | 0.31 | 0.11 | 0.511 | 7.36% |
| P08 | 1.888 | 1.462 | 0.426 | 0.186 | 0.667 | 5.1% |
| S10 | 2.1 | 1.406 | 0.695 | 0.495 | 0.895 | 7.38% |
| U01 | 2.358 | 2.027 | 0.331 | 0.2 | 0.461 | 17.2% |
| U02 | 2.288 | 2.07 | 0.218 | 0.07 | 0.367 | 13.36% |
| U03 | 2.46 | 2.395 | 0.064 | -0.055 | 0.184 | 20.69% |
| **Meta analysis** | | | **0.258** | **0.204** | **0.312** | |

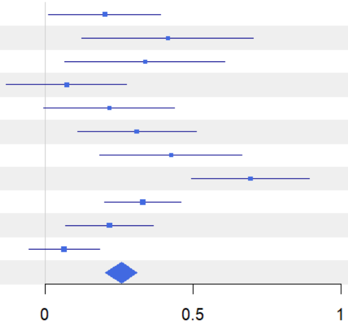


**Figure S29**: Forest plot for functional gamma diversity of order q = 2 showing results of coverage-standardised comparisons between ESBC and C-districts of each individual experimental landscape and across landscapes.

**Alpha with Coverage = 0.9**

| regionSite | FD(q=2) E | FD(q=2) C | Difference | LCL | UCL | W(fixed) |
|---|---|---|---|---|---|---|
| B04 | 1.813 | 1.322 | 0.491 | 0.285 | 0.696 | 3.88% |
| B05 | 1.728 | 1.351 | 0.377 | 0.179 | 0.575 | 4.18% |
| B06 | 1.793 | 1.068 | 0.725 | 0.463 | 0.986 | 2.39% |
| B07 | 1.764 | 1.268 | 0.496 | 0.301 | 0.691 | 4.31% |
| H09 | 1.475 | 1.242 | 0.234 | 0.019 | 0.448 | 3.56% |
| L11 | 1.054 | 1.386 | -0.333 | -0.465 | -0.201 | 9.4% |
| P08 | 1.647 | 1.358 | 0.288 | 0.083 | 0.493 | 3.9% |
| S10 | 1.542 | 1.272 | 0.27 | 0.126 | 0.414 | 7.88% |
| U01 | 2.216 | 1.694 | 0.522 | 0.436 | 0.609 | 21.68% |
| U02 | 2.177 | 1.597 | 0.58 | 0.485 | 0.676 | 17.97% |
| U03 | 2.372 | 2.353 | 0.019 | -0.07 | 0.108 | 20.85% |
| **Meta analysis** | | | **0.305** | **0.264** | **0.345** | |

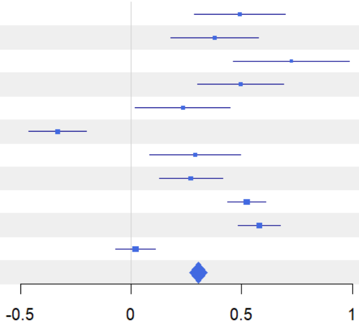


**Figure S30**: Forest plot for functional alpha diversity of order q = 2 showing results of coverage-standardised comparisons between ESBC and C-districts of each individual experimental landscape and across landscapes.

**1-S with Coverage = 0.9**

| regionSite | FD(q=2) E | FD(q=2) C | Difference | LCL | UCL | W(fixed) |
|---|---|---|---|---|---|---|
| B04 | 0.049 | 0.178 | -0.129 | -0.218 | -0.039 | 2.67% |
| B05 | 0.134 | 0.145 | -0.011 | -0.107 | 0.086 | 2.29% |
| B06 | 0.179 | 0.429 | -0.249 | -0.346 | -0.153 | 2.28% |
| B07 | 0.025 | 0.289 | -0.264 | -0.347 | -0.182 | 3.11% |
| H09 | 0.041 | 0.041 | 0 | -0.105 | 0.105 | 1.95% |
| L11 | 0.499 | 0.072 | 0.427 | 0.358 | 0.495 | 4.58% |
| P08 | 0.17 | 0.091 | 0.08 | -0.023 | 0.183 | 2.02% |
| S10 | 0.303 | 0.149 | 0.154 | 0.06 | 0.248 | 2.43% |
| U01 | 0.042 | 0.177 | -0.135 | -0.165 | -0.105 | 23.24% |
| U02 | 0.031 | 0.249 | -0.218 | -0.25 | -0.186 | 21.19% |
| U03 | 0.025 | 0.021 | 0.004 | -0.021 | 0.029 | 34.25% |
| **Meta analysis** | | | **-0.069** | **-0.084** | **-0.054** | |

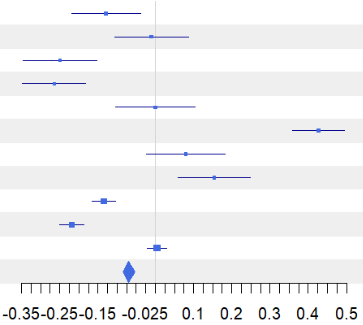


**Figure S31**: Forest plot for functional beta (1-S) diversity of order q = 2 showing results of coverage-standardised comparisons between ESBC and C-districts of each individual experimental landscape and across landscapes.

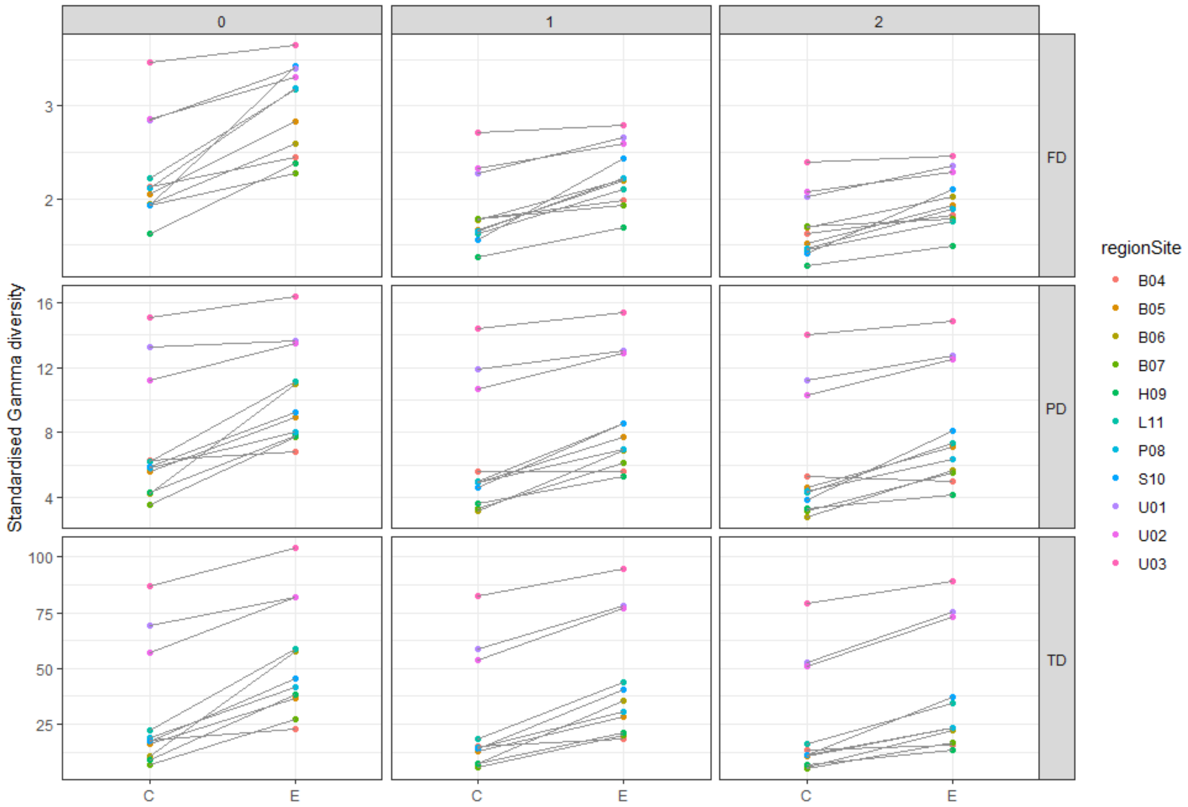


**Fig. S32**: Standardised gamma diversity at a sample coverage of 0.9 across all three diversity facets—taxonomic diversity (TD), phylogenetic diversity (PD), and functional diversity (FD)—for each district. Points represent individual districts (C = control, E = ESBC), with grey lines connecting paired control and ESBC districts.

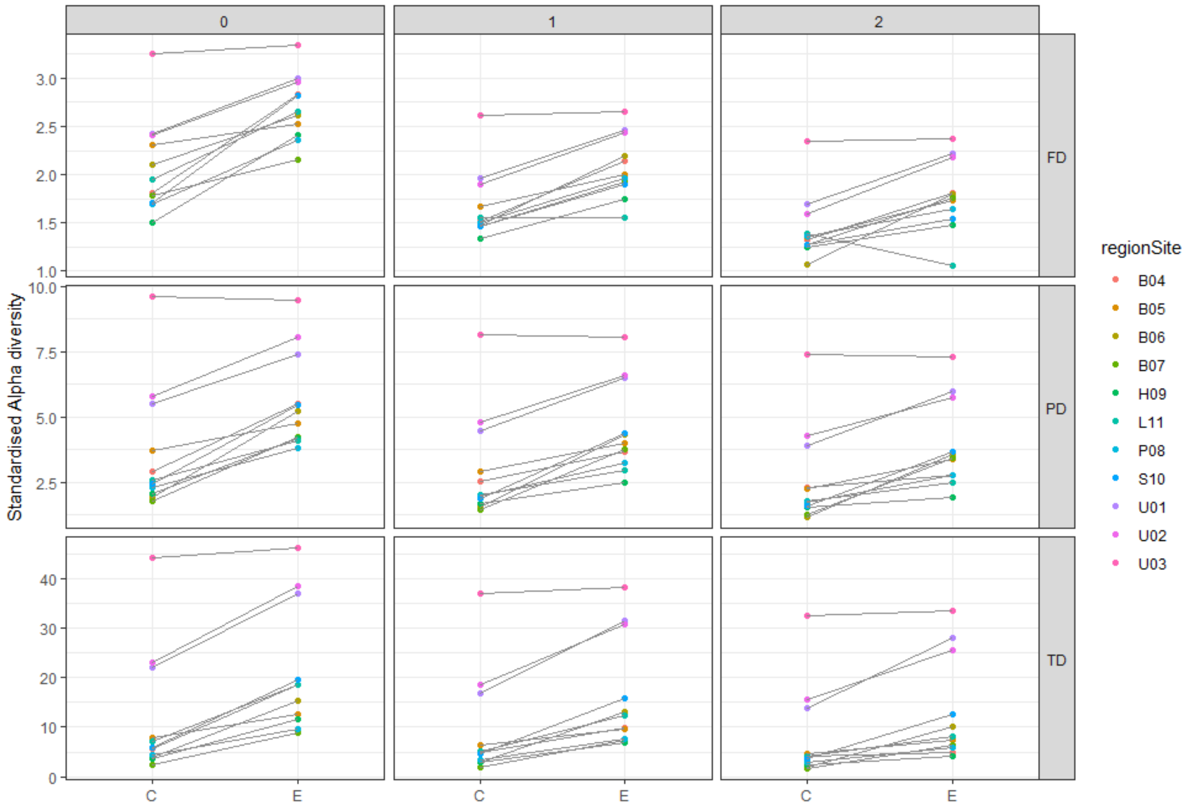


**Fig. S33**: Standardised alpha diversity at a sample coverage of 0.9 across all three diversity facets—taxonomic diversity (TD), phylogenetic diversity (PD), and functional diversity (FD)—for each district. Points represent individual districts (C = control, E = ESBC), with grey lines connecting paired control and ESBC districts.

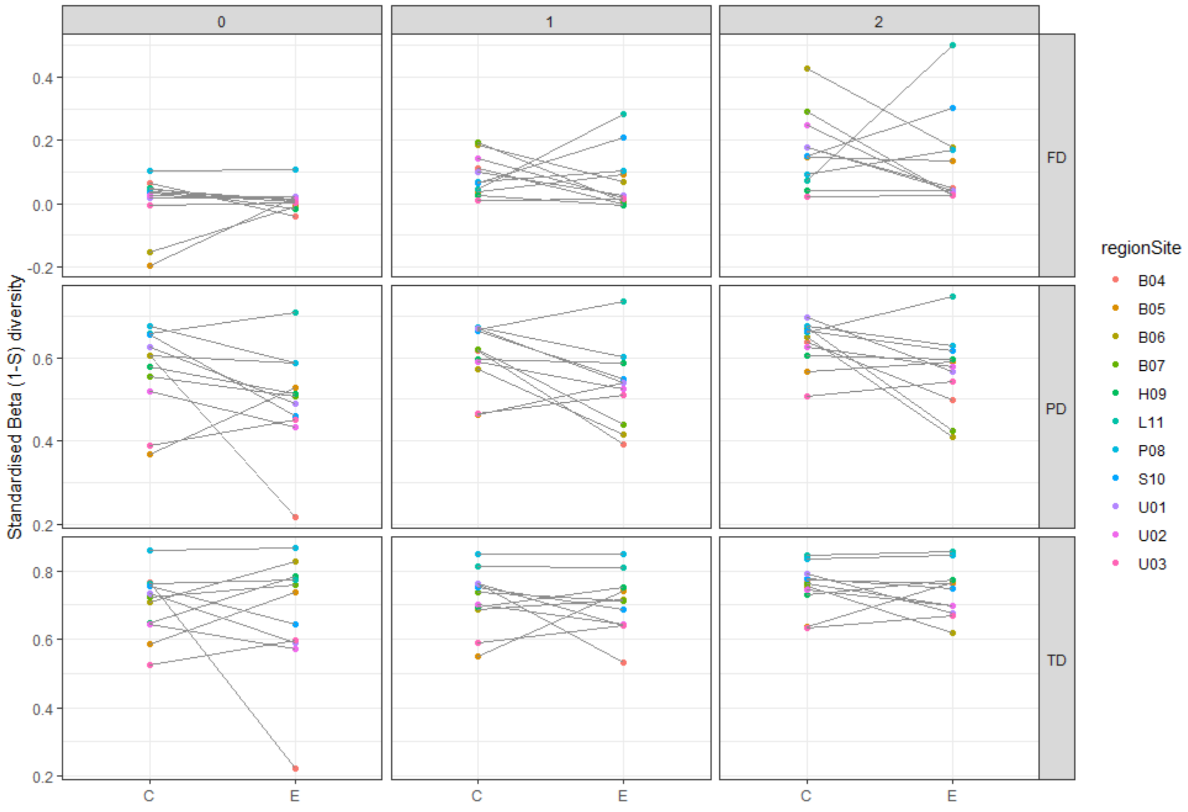


**Fig. S35**: Standardised beta diversity (1 − S) at a sample coverage of 0.9 across all three diversity facets—taxonomic diversity (TD), phylogenetic diversity (PD), and functional diversity (FD)—for each district. Points represent individual districts (C = control, E = ESBC), with grey lines connecting paired control and ESBC districts.

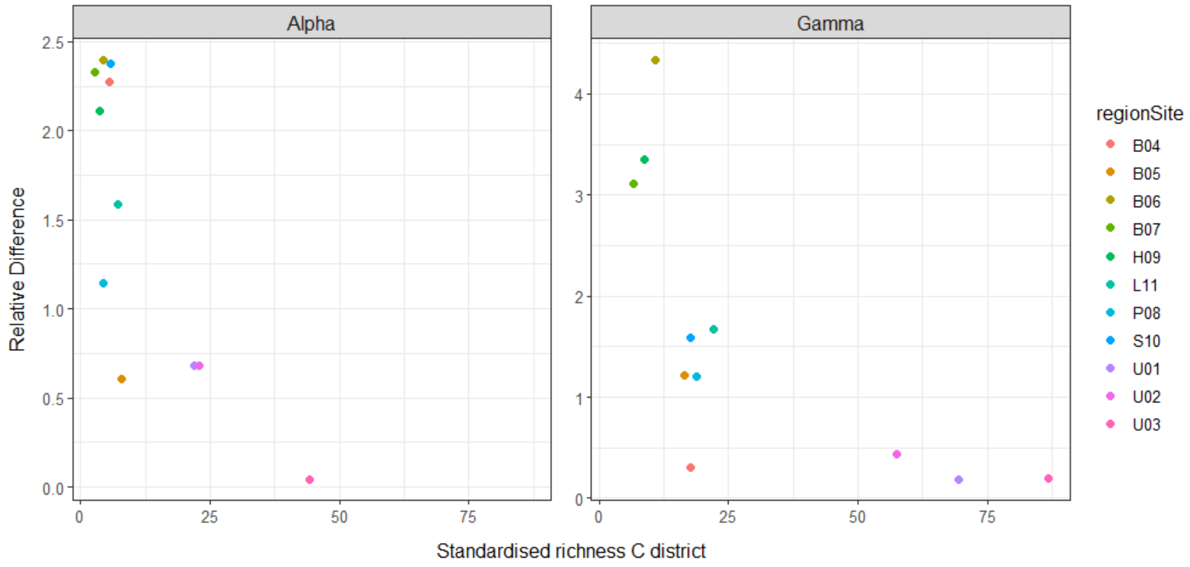


**Figure S35**: Standardised taxonomic gamma diversity (q = 0) at a coverage of 0.9 of each control district plotted against the relative difference compared to the ESBC district. A relative difference of 1 corresponds to a 100% increase in diversity.

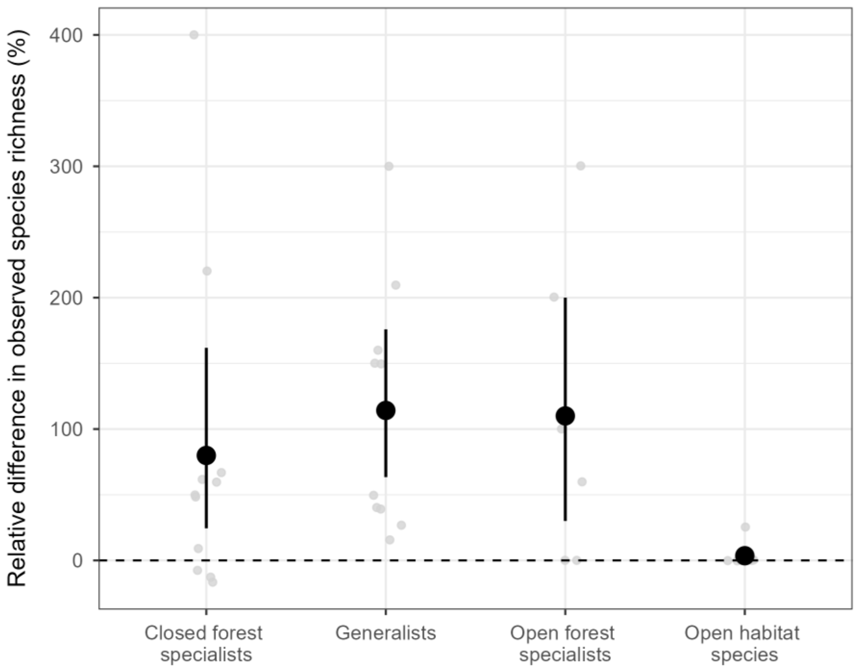


**Figure S36**: Effects of an enhancement of structural heterogeneity in forest landscapes on species' affinity to forests**.** Light grey points represent the relative differences in % in the observed species richness per forest affinity category between the ESBC and control district across experimental landscapes, and error bars denote the 95%-confidence intervals computed by bootstrapping. Relative differences were calculated by dividing the observed difference in species richness by the control richness per category and site. Note than no relative increase could be calculated for those sites where observed richness of a category was zero. Species are categorised as closed forest specialists, open forest specialists, generalists and open habitat species. Positive differences denote higher observed species richness in ESBC districts, while negative differences denote the opposite. Note, that a difference of zero (in this dataset) indicates that the category was absent in both districts.
